# Engineering mtDNA Deletions by Reconstituting End-Joining in Human Mitochondria

**DOI:** 10.1101/2024.10.15.618543

**Authors:** Yi Fu, Max Land, Ruobing Cui, Tamar Kavlashvili, Minsoo Kim, Toby Lieber, Keun Woo Ryu, Emily DeBitetto, Ignas Masilionis, Rahul Saha, Meril Takizawa, Daphne Baker, Marco Tigano, Ed Reznik, Roshan Sharma, Ronan Chaligne, Craig B. Thompson, Dana Pe’er, Agnel Sfeir

## Abstract

Recent breakthroughs in the genetic manipulation of mitochondrial DNA (mtDNA) have enabled the precise introduction of base substitutions and the effective removal of genomes carrying harmful mutations. However, the reconstitution of mtDNA deletions responsible for severe mitochondrial myopathies and age-related diseases has not yet been achieved in human cells. Here, we developed a method to engineer specific mtDNA deletions in human cells by co-expressing end-joining (EJ) machinery and targeted endonucleases. As a proof-of-concept, we used mito-EJ and mito-ScaI to generate a panel of clonal cell lines harboring a ∼3.5 kb mtDNA deletion with the full spectrum of heteroplasmy. Investigating these isogenic cells revealed a critical threshold of ∼75% deleted genomes, beyond which cells exhibited depletion of OXPHOS proteins, severe metabolic disruption, and impaired growth in galactose-containing media. Single-cell multiomic analysis revealed two distinct patterns of nuclear gene deregulation in response to mtDNA deletion accumulation; one triggered at the deletion threshold and another progressively responding to increasing heteroplasmy. In summary, the co-expression of mito-EJ and programable nucleases provides a powerful tool to model disease-associated mtDNA deletions in different cell types. Establishing a panel of cell lines with a large-scale deletion at varying levels of heteroplasmy is a valuable resource for understanding the impact of mtDNA deletions on diseases and guiding the development of potential therapeutic strategies.

**Graphical Abstract:** 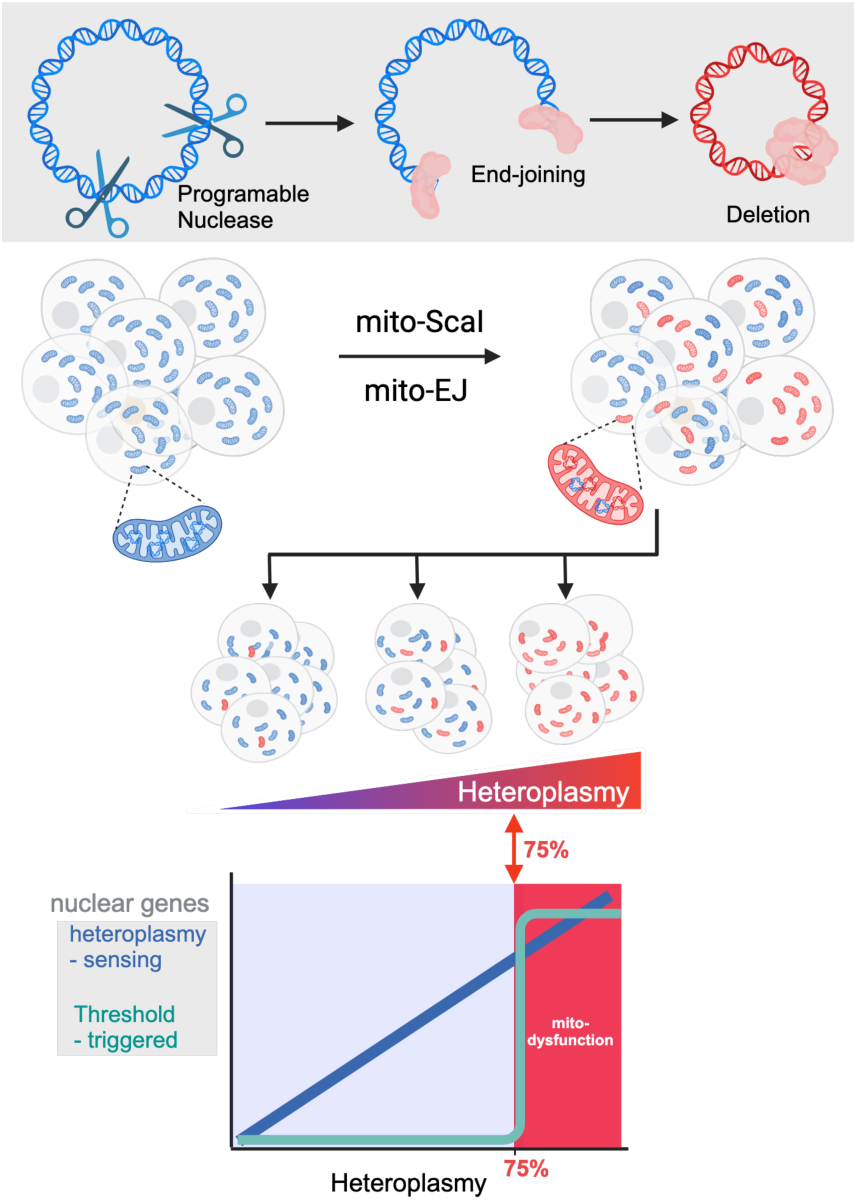

**Highlights:**

- Combining prokaryotic end-joining with targeted endonucleases generates specific mtDNA deletions in human cells
- Engineering a panel of cell lines with a large-scale deletion that spans the full spectrum of heteroplasmy
- 75% heteroplasmy is the threshold that triggers mitochondrial and cellular dysfunction
- Two distinct nuclear transcriptional programs in response to mtDNA deletions: threshold-triggered and heteroplasmy-sensing

## INTRODUCTION

Mitochondria possess their own genome, a circular DNA that encodes subunits of the oxidative phosphorylation (OXPHOS) complexes responsible for cellular respiration and ATP production. Mitochondrial DNA (mtDNA) is prone to mutations, including base substitutions and deletions, often leading to mitochondrial diseases. Large-scale mtDNA deletions, ranging from 1.8 to 8 kilobases (kb), are a common cause of mitochondrial pathologies, including Progressive External Ophthalmoplegia (PEO), Pearson syndrome, and Kearns-Sayre syndrome (KSS) and are also associated with aging and neurodegeneration ^1–3^. mtDNA deletions are predominantly sporadic, not typically inherited, and accumulate over time, particularly in post-mitotic tissues such as muscles and the brain ^4,5^. In some cases, the accumulation of mtDNA deletions is associated with mutations in nuclear-encoded mitochondrial replisome components, including PolG and Twinkle ^6–8^. Typically, mutant mitochondrial genomes co-exist with wild-type mtDNA in a state of heteroplasmy and manifest clinically as heteroplasmy surpasses a critical threshold ^9^. Due to the difficulty in editing mtDNA and modeling disease-causing mtDNA deletions in human cells, the mechanisms underlying their pathogenicity remain unclear. Particularly, how deleted genomes impair mitochondrial and cellular function, the threshold at which they manifest pathologically, and what transcriptional response they activate are unknown.

The dynamics and functional consequences of mtDNA deletions have been primarily studied in model organisms, namely worms and flies. In *C. elegans*, a 3.1 kb deletion caused by mutagenesis initiates a mitochondrial stress response that facilitates mtDNA replication and maintenance across generations ^10–12^. Conversely, in *D. melanogaster*, mtDNA deletions produced by the co-expression of a restriction enzyme and T4 DNA ligase are eliminated from muscle cells through autophagy and activating the PINK1/parkin pathway^13^. Several mouse models of mtDNA deletions have been developed, including transmitochondrial “Mito-mice” and the Twinkle transgenic mouse, which successfully replicate features of mitochondrial myopathy^6,14,15^. However, interpreting results from these mouse models is complicated by concurrent deletions and a reduced mtDNA copy number. Studies on human mtDNA deletions have primarily been limited to correlation analyses of patient tissue biopsies, with few cell models available ^16^. For instance, single-cell multiomics of blood samples from patients carrying mtDNA deletions identified patterns consistent with purifying selection associated with rapidly dividing T cells ^17^. Available cell-based models of mtDNA deletions are limited to cybrids, which often exhibit mismatches between nuclear and mitochondrial genomes, and induced pluripotent stem cells (iPSCs) derived from patient cells^18–20^. These models are frequently constrained by the limited types of deletions and levels of heteroplasmy they harbor. To investigate the mtDNA deletion dynamics in human cells, probe the tissue specificity underlying its disease manifestation, and explore potential treatment for associated mitochondrial myopathies, it is necessary to develop tools capable of modeling mtDNA deletions at defined heteroplasmy levels in a cell-type-specific manner.

Editing mtDNA has been particularly challenging due to multiple copies of the circular genome per cell and the inability to deliver nucleic acids across the mitochondrial inner membrane. However, recent innovations based on programmable nucleases, including transcription activator-like effectors nucleases (TALENs), zinc fingers nucleases, and restriction enzymes, have seen success in inducing DNA breaks in mtDNA that lead to heteroplasmic shifts in favor of intact genomes ^21–23^. Mitochondrial-targeted transcription activator-like effectors (TALEs) fused to bacterial deaminases successfully modeled disease-associated point mutations in mtDNA^24^ and transformed the study of mtDNA base substitutions. However, the progress in developing tools to model the highly toxic deletions in mtDNA lags. A major obstacle to generating mtDNA deletions is the lack of double-strand break (DSB) repair machinery in the mitochondria, where broken DNA ends are degraded rather than joined to form deletions ^25–27^.

In this study, we employed a cross-kingdom approach to devise a simple, efficient, and tractable two-component system to engineer specific mtDNA deletions in different cell types. Our method involves co-expressing minimal DNA end-joining (EJ) machinery from Mycobacterium or T4 bacteriophage with site-specific mitochondrial-targeted nucleases in human cells. To validate this method, we co-expressed mito-EJ and mito-ScaI in epithelial cells, successfully engineering a panel of cells with a ∼3.5 kb mtDNA deletion (mtDNA^ΔScaI^) comprising the full spectrum of heteroplasmy levels. In-depth characterization of cells with different mtDNA^ΔScaI^ abundance identified a critical threshold of ∼75% heteroplasmy. Beyond this threshold, cells had impaired OXPHOS, lost their ability to engage in the tricarboxylic acid (TCA) cycle, and displayed growth defects when cultured in galactose-rich media. We applied an improved single-cell DOGMAseq^28^ on cells harboring mtDNA^ΔScaI^ and detected two distinct gene expression programs. A set of nuclear genes were deregulated in cells that surpassed the critical heteroplasmy threshold. A distinct and previously unappreciated group of heteroplasmy-sensing genes displayed a gradual response, whereby their deregulation correlated with the abundance of mtDNA^ΔScaI^. Finally, we demonstrated that our deletion-generating method can induce various mtDNA deletions in somatic cells and induced pluripotent stem cells (iPSCs). In summary, combining mito-EJ with programable nucleases offers a genetic tool to model disease-associated mtDNA deletions. The panel of isogenic cells with the gradient of mtDNA^ΔScaI^ heteroplasmies provides a unique resource for investigating the pathogenicity of deletions and testing treatment strategies for numerous mitochondrial diseases.

## RESULTS

### Reconstituting end-joining (EJ) in human mitochondria leads to the accumulation of mtDNA deletions

Mammalian mitochondria lack the machinery for repairing DNA double-strand breaks (DSBs). As a result, cleaved mitochondrial genomes are rapidly degraded and only rarely get rejoined, forming mtDNA deletions ^25–27^. To trigger mtDNA deletion formation, we expressed the non-homologous end-joining (NHEJ) machinery from Mycobacterium tuberculosis in human mitochondria (Figure S1A). The mitochondrial end-joining (mito-EJ) system comprised two components, the DNA-binding protein, Ku, and Ligase D (LigD) ^29^. Independently, we explored whether a bacteriophage-derived T4 DNA ligase could effectively ligate broken mtDNA ends. Ku, LigD, and T4 ligase, each codon-optimized and amended with a mitochondrial localization signal (MLS), were stably expressed in human retinal pigmental epithelial (ARPE-19) cells, previously shown to support the accumulation of mtDNA mutations^30^. Immunofluorescence and subcellular fractionation followed by Western blot confirmed proper mitochondrial targeting of mito-EJ components (Figure S1B-E). We first assessed the functionality of the cross-kingdom mito-EJ approach by expressing a mitochondria-targeted restriction enzyme ApaLI (mito-ApaLI) ^31^, which creates a single mitochondrial DSB. Southern blot analysis revealed that the degradation of linear mtDNA was delayed in cells expressing mito-Ku, suggesting a protective role for Ku in shielding mtDNA from degradation (Figure S1F). To detect ligation activity by LigD, we expressed a mitochondria-targeted ScaI (mito-ScaI)^32^, which cleaves five sites in the major arc of human mtDNA. The distal ScaI sites flank a 3.5 kb mtDNA fragment that overlaps with seven protein-coding genes and three tRNAs, commonly excised in large-scale disease-associated mtDNA deletions ^33^ (Figure 1A and S1G). We designed primers that captured the joining of the most distant ScaI sites (Figure S1G). Using quantitative PCR (qPCR), we detected a significant accumulation of mtDNA^ΔScaI^ deletions in cells expressing Ku and LigD or T4 ligase alone (Figure 1B and S1H-J). In addition, we detected smaller amplicons due to DNA end-resection beyond the most distal ScaI sites (Figure S1F and S1I-J).

**Figure 1.**
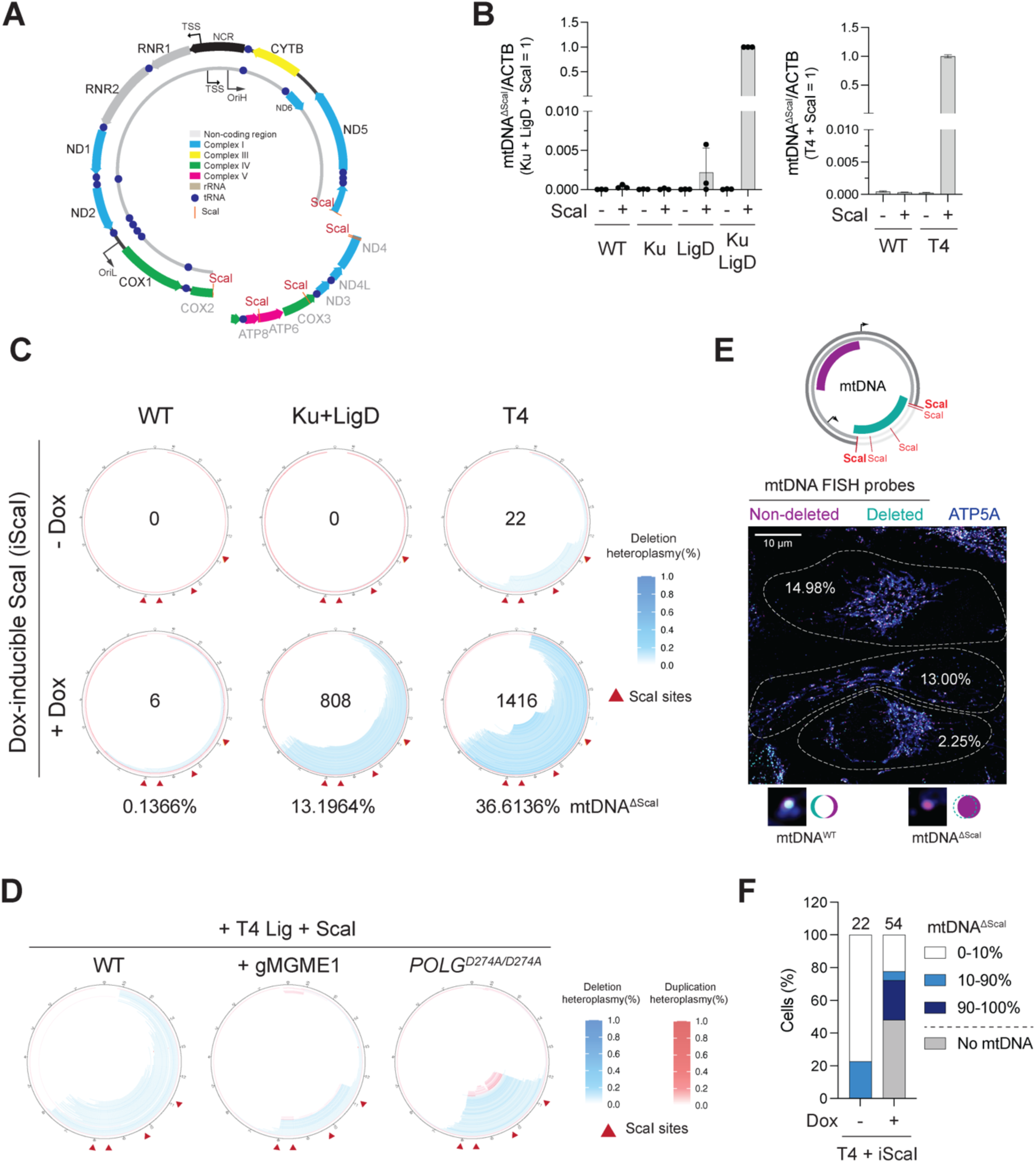
Reconstitution of end-joining in the mitochondria results in the accumulation of mtDNA deletions. **(A)** Human mtDNA is a 16.5 kb circular genome. The heavy strand is transcribed into a polycistronic transcript, which is then cleaved into individual tRNA, rRNA, and mRNAs coding for the indicated subunits of the OXPHOS complex. The most distal ScaI sites flank a 3.5 kb region that overlaps with seven peptides and three tRNA genes. (B) Quantitative PCR (qPCR) analysis of the levels of mtDNA deletions in ARPE-19 cells that stably expressed mitochondria end-joining (mito-EJ) and were transfected with mito-ScaI. Cells were analyzed four days post-transfection. The presence of mtDNA^DScaI^ was detected using primers flanking ScaI sites and normalized to nuclear DNA locus ACTB. Left panel, values were normalized to Ku + LigD cells transfected with mito-ScaI. Right panel, values were normalized to T4 ligase-expressing cells transfected with mito-ScaI. Values are mean ± s.d. from *n* = 3 technical replicates. **(C)** Circular plots depict mtDNA deletions (blue) identified in the indicated samples. ARPE-19 cells expressing mito-EJ and doxycycline (Dox) inducible mito-ScaI were treated with Dox for 12 hours and harvested on Day 5 for ATAC-seq. The number of unique deletions is highlighted at the center of each circular plot, with the overall percentage of mtDNA^ΔScaI^ indicated below. Table S4 lists all detected deletions. **(D)** Circular plots displaying deleted (blue) and duplicated (red) mtDNA sequences in cells with the indicated genotype and co-expressing mito-ScaI and mitochondrial-targeted T4 ligase. All deletions are listed in Table S1. **(E)** Fluorescent *in situ* hybridization (FISH) staining on mtDNA in ARPE-19 cells expressing mito-EJ and Dox-inducible mito-ScaI. Cells were treated with Dox for 24 hours and analyzed on Day 5. The schematic highlights recognition sites of two independent FISH probes that distinguish intact mtDNA (pink) from total mtDNA molecules (green). Overlapping signals from both probes distinguish mtDNA^WT^ from mtDNA^ΔScaI^. ATP5A staining in blue marks mitochondria. Highlighted are three cells and the percentage of mtDNA^ΔScaI^ in each cell is noted. **(F)** Graph displaying the percentage of cells with the indicated mtDNA^ΔScaI^ heteroplasmy quantified by mtDNA FISH. The total number of analyzed cells is highlighted in the graph.

To further characterize mtDNA in cells co-expressing mito-EJ and doxycycline-inducible mito-ScaI, we performed transposase-accessible chromatin with sequencing (ATAC-seq). We applied the MitoSAlt computational pipeline that detects mtDNA deletions ^34^. While ScaI-cleavage of mtDNA in control cells resulted in minimal deletion accumulation, cells with reconstituted mito-EJ showed significant enrichment of deleted genomes. We noted the accumulation of deletions extending beyond the distal ScaI sites (Figure 1C and S2A and Table S1), suggesting competition between the DNA end-joining and the degradation of cleaved DNA by exonucleases. Consistent with the previously documented role of *POLG* and *MGME1* in degrading linear mtDNA^25,27^, extended deletions were diminished upon co-expression of mito-ScaI and mito-EJ in cells lacking *MGME1* or carrying an exonuclease-dead allele of *POLG.* This resulted in a decrease in overall deletion heterogeneity. Furthermore, mitochondrial DSB repair events in *POLG^D274A/D274A^* and *MGME1^-/-^* cells often resulted in small duplications, potentially due to the joining of unresected fragments with the mtDNA backbone (Figure 1D and S2B-D and Table S1)^35^.

To determine the heterogeneity of mtDNA deletions in individual cells, we performed fluorescence in situ hybridization (FISH) using two probes. One probe hybridized to the deleted region, allowing for the detection of wild-type mtDNA, while a second probe recognized all mitochondrial genomes (Figure 1E and S3A). Co-staining with both probes highlighted a fraction of cells comprising deletions at different heteroplasmy levels (Figure 1E-F). These results confirmed that implementing mito-EJ in retinal epithelial cells resulted in the accumulation of mtDNA^ΔScaI^ with cell-to-cell heterogeneity in heteroplasmy levels.

### Establishing isogenic cells with the full spectrum of mtDNA**^ΔScaI^** heteroplasmy

We leveraged mtDNA^ΔScaI^ heterogeneity in the population of ARPE-19 cells expressing mito-T4 and mito-ScaI to isolate single-cell clones and establish cell lines carrying the entire range of heteroplasmy. Genotyping PCR revealed that most isolated clones exhibited a dominant mtDNA deletion (Figure S3A-D). We quantified the percentage of mtDNA^ΔScaI^ molecules in the clonally derived cell lines using qPCR and FISH. Our results suggested that the average heteroplasmy level in most cell lines had a narrow distribution that remained stable over time (Figure 2A and S3E-G). We selected twelve representative clones spanning the mtDNA^ΔScaI^ heteroplasmy spectrum for in-depth analyses of mitochondrial and metabolic functions (Figure 2A). The mtDNA deletions from ScaI cleavage extended across the major arc of the mitochondrial genome and mimicked large-scale deletion associated with mitochondrial diseases ^36^. Specifically, mtDNA^ΔScaI^ genomes lack at least five protein-coding genes (ATP8, ATP6, COX3, ND3, and ND4L, with COX2 and ND4 being partially deleted) and three mitochondrial tRNAs (MT-TK, MT-TG, MT-TR) (Figure 1A). We performed Western blot analyses to assess the impact of mtDNA deletion accumulation on the abundance of mitochondrial and nuclear-encoded OXPHOS subunits. Our results revealed that all mitochondrial encoded OXPHOS proteins, including those expressed from regions retained in mtDNA^ΔScaI^, were depleted in cells with ≥75% heteroplasmy (Figure 2B). Additionally, we observed a simultaneous reduction in the nuclear-encoded subunits of respiratory complexes, with the exception of ATP5A (a nuclear-encoded Complex V subunit). Notably, other mitochondrial-resident proteins, such as transcription factor A, mitochondrial (TFAM), and voltage-dependent anion channel 1 (VDAC1), maintained stable expression levels (Figures 2B).

**Figure 2.**
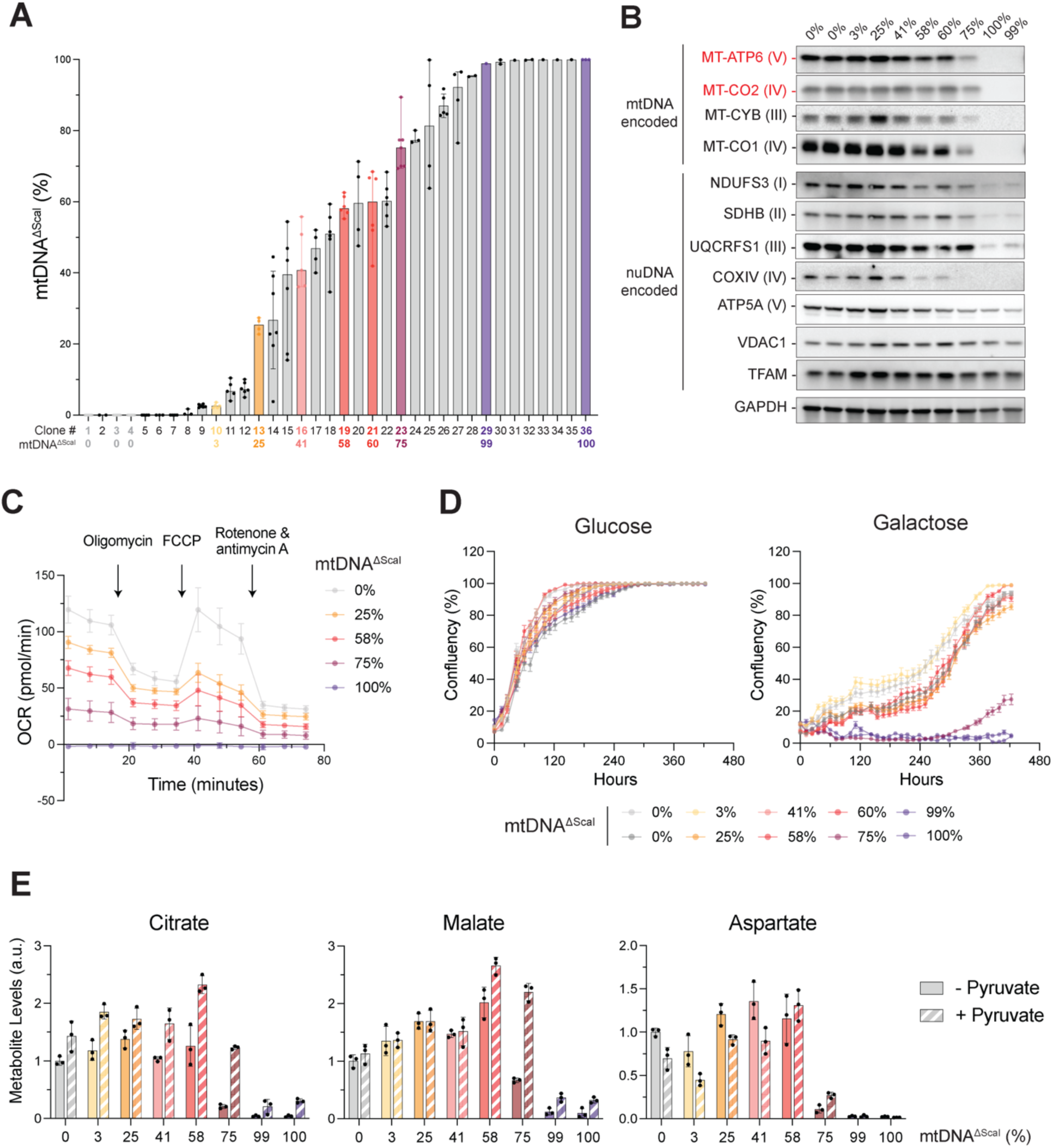
Establishing isogenic cells with a range of mtDNA^ΔScaI^heteroplasmy. **(A)** Representative clonal cells isolated from a population of ARPE-19 cells expressing T4 ligase and Dox-inducible mito-ScaI. The mtDNA^ΔScaI^ heteroplasmy level in each clone was quantified by qPCR. Representative clones highlighted in color were prioritized in the study. Plotted values are median ± s.d. from *n* = 1 - 6 technical replicates. **(B)** Protein levels of the subunits of OXPHOS complexes, VDAC1, and TFAM in mtDNA^ΔScaI^ clones with the indicated heteroplasmy. mtDNA-encoded subunits deleted in mtDNA^ΔScaI^ are highlighted in red. **(C)** Analysis of Oxygen Consumption Rate (OCR) in the indicated mtDNA^ΔScaI^ clones using Seahorse assay. **(D)** Growth curves of indicated mtDNA^ΔScaI^ clones in 25 mM glucose (left) and 10 mM galactose (right) media supplemented with 1mM pyruvate and 50μg/ml uridine. **(E)** TCA metabolites in the indicated mtDNA^ΔScaI^ clones cultured in glucose media with or without pyruvate were measured by gas chromatography-mass spectrometry (GC-MS). Values represent mean ± s.d. from *n* = 3 technical replicates.

Next, we conducted a Seahorse analysis to determine the effect of increasing heteroplasmy on oxygen consumption rate (OCR). Analysis of representative cell lines demonstrated a progressive decline in basal and maximum respiration in cells with ≥25% mtDNA^ΔScaI^ heteroplasmy (Figure 2C). Cells with heteroplasmy levels exceeding 75% displayed severe respiration defects and abnormal mitochondrial morphology (Figure S4A). Despite the profound impairment in oxygen consumption in cells with high mtDNA^ΔScaI^ heteroplasmy, all clonally derived cells proliferated normally in glucose-rich media supplemented with pyruvate, uridine, and nonessential amino acids (NEAA) (Figure 2D) and retained the ability to maintain a mitochondrial membrane potential (Figure S4B). In contrast, cells with ≥75% heteroplasmy exhibited severe growth defects when glucose-rich media was replaced with galactose, and clones near homoplasmy underwent apoptosis (Figure 2D and S4C). These results confirmed that retinal epithelial cells can tolerate high levels of mtDNA^ΔScaI^ in glucose-rich media by relying primarily on glycolysis for ATP synthesis ^37^. In galactose media, where cells depend on OXPHOS for ATP production, defects resulting from mtDNA deletions become evident. Exposing cells to metabolic stress by replacing glucose with galactose resulted in a slight decrease in mtDNA^ΔScaI^ heteroplasmy in most cells (Figure S2E). In conclusion, the reconstitution of mito-EJ in the presence of mito-ScaI enabled us to establish a panel of isogenic cell lines with heteroplasmy ranging from 0 to 100% as a valuable resource to investigate the metabolic and cellular dysfunction in response to mtDNA deletions in human cells.

### A critical metabolic threshold of 75% heteroplasmy for mtDNA^ΔScaI^

Growth analysis of clonally derived cells comprising a gradient of mtDNA^ΔscaI^ revealed a threshold of 75% heteroplasmy, beyond which cells lose OXPHOS components and exhibit mitochondrial and cellular defects. A functional electron transport chain (ETC) couples the oxidative process (oxidizing NADH to generate NAD+) with phosphorylation (ATP generation). NAD+ is an essential electron acceptor for various metabolic pathways, including the TCA cycle and glycolysis. Deficiency of NAD+ due to ETC inhibition hinders the TCA cycle and cell proliferation, rendering cells auxotrophic for exogenous electron acceptors^38^. Pyruvate is a commonly used electron acceptor, as reducing pyruvate via lactate dehydrogenase (LDH) oxidizes NADH to regenerate NAD+. The electron acceptor auxotrophy has been observed in ρ0 cells and homoplasmic cybrid cell lines and was also evident in cells treated with ETC inhibitors^38–40^. However, the exact threshold of defective mitochondrial genomes that trigger such metabolic dependency has not been determined.

We cultured cells with the full spectrum of heteroplasmy in high glucose without supplementing the media with exogenous pyruvate and noted significant growth defects with ≥ 75% heteroplasmy (Figure S4D-E). We observed a concomitant loss of TCA metabolites, including citrate and malate (Figure 2E). It has been well-established that a major role of mitochondrial respiration is to support aspartate synthesis, and aspartate deficiency restricts cell proliferation upon electron acceptor deficiency^39,40^. Aspartate levels were significantly reduced in cells with 75% heteroplasmy and almost completely depleted in cells with over 90% mtDNA^ΔScaI^ (Figure 2E). Supplementing the media with exogenous pyruvate rescued cell growth and partially restored TCA metabolites in cells with 75% of mtDNA^ΔScaI^ (Figure 2E and Figure S4D-E). However, TCA cycle metabolites and aspartate levels remained critically low in homoplasmic cells despite pyruvate supplementation. These findings demonstrate that while multi-copy mitochondrial genomes offer a metabolic buffer, ARPE-19 cells experience a deficiency in electron acceptors and a disruption in redox balance when mtDNA deletions surpass a 75% heteroplasmy threshold.

### Single-cell multiomics uncover the dynamics of mitochondrial RNA as a function of heteroplasmy

So far, studies have predominantly focused on probing the cellular response to large-scale mtDNA deletions only after they manifest in disease states, thus capturing merely the end-stage effects. We sought to harness the full spectrum of mtDNA^ΔScaI^ heteroplasmy to systematically record the dynamic transcriptional network across the continuum of mtDNA deletion levels and in single cells. Given the dynamic nature and variability of mtDNA heteroplasmy, even within clones, we applied an optimized DOGMAseq protocol (high LLL DOGMAseq) with enhanced mtDNA recovery. With this approach, we analyzed mtDNA based on single-cell ATAC-seq (scATAC-seq) and transcripts through single-cell RNA-seq (scRNA-seq)^28,41^. After pre-processing the scATAC-seq data, we estimated heteroplasmy in single cells by computing the ratio of total read coverage from the deleted region (8013-11460 bp) to an undeleted region (1231-4678 bp) (Figure S5A) (see methods section). To validate the heteroplasmy estimation method, we first conducted a pilot experiment involving a set of eight clones with predetermined heteroplasmy (Figure 2A and S3G). Analyses of the average deletion coverage and per-cell heteroplasmy distribution in each clone were consistent with the results obtained using qPCR and FISH analysis (Figure S5A-B and S3G).

Having validated the computational pipeline for estimating heteroplasmy, we explored the effects of mtDNA^ΔScaI^ levels on the dynamics of the transcriptome, initially focusing on RNA encoded by the mitochondria (mtRNA). We pooled together an equal number of cells from 46 mtDNA^ΔScaI^ clones harboring the full spectrum of heteroplasmy. We treated the pooled cells for three days in three different culture conditions: 1) Glucose + Pyruvate, 2) Glucose – Pyruvate, and 3) Galactose + Pyruvate (Figure 3A). While Glucose + Pyruvate supports normal growth across all heteroplasmy levels, media that either lack pyruvate or contain galactose challenge the ETC differently. The absence of pyruvate affects the ETC’s ability to maintain redox balance, whereas the presence of galactose limits its capacity to generate ATP, both in a manner dependent on heteroplasmy. We performed DOGMAseq and concomitantly determined mtDNA heteroplasmy and mtRNA levels per cell (Figure 3A). We categorized single cells into bins based on their heteroplasmy levels: wild-type (WT; 0 – 0.5%), low (0.5% – 30%), medium (30% – 65%), and high (65% – 100%) heteroplasmy (Figure 3B, Figure S6A-B). Each category, except for high heteroplasmy galactose, contained a sufficient number of cells (> 690) to ensure the robustness of our conclusions. Consistent with growth curve analysis, cells with high heteroplasmy dropped out in galactose medium in three days (Figure 3B). We then measured the average expression of mitochondrial mRNA from the scRNA-seq data (Figure 3C). As expected, transcripts encoded by genes within the deleted region were depleted at high heteroplasmy. Conversely, mitochondrial-encoded genes outside the deleted region, namely ND1 and ND2, were significantly upregulated at high heteroplasmy in cells growing in glucose-rich media (Figure 3C).

**Figure 3.**
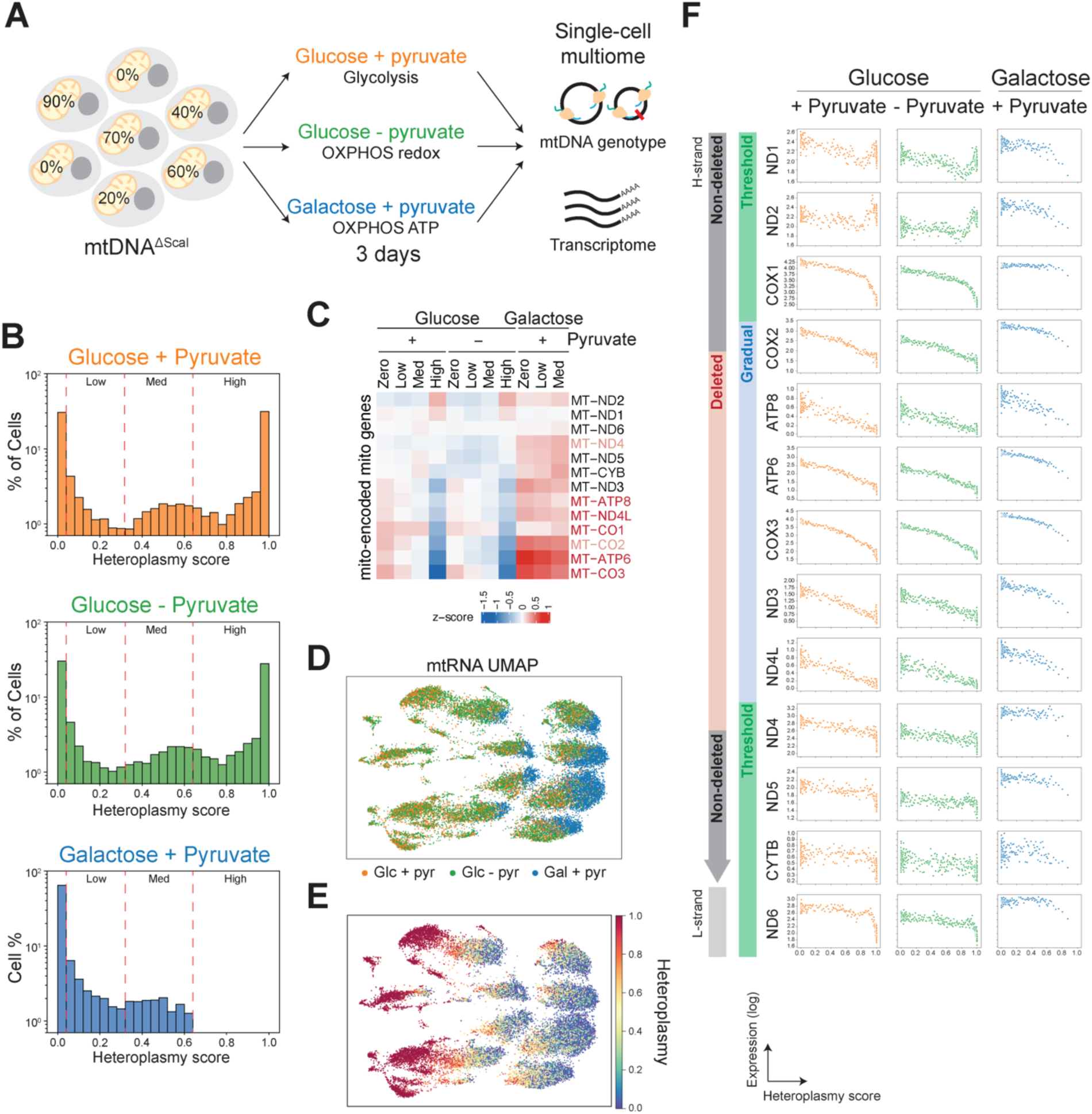
Single-cell multiome analysis reveals the dynamics of mitochondrial transcripts as a function of heteroplasmy. **(A)** Schematic of the experimental outline. 46 clonal cell populations with heteroplasmy levels ranging from 0-100% were mixed in equal cell numbers and cultured for three days in media containing Glucose + Pyruvate, Glucose – Pyruvate, and Galactose. and analyzed by modified DOGMAseq (see Methods section for description). **(B)** Graph representing the percentage of cells with each distribution of heteroplasmy levels in single cells in the indicated treatment, captured by scATAC-seq in DOGMAseq. Cells were categorized according to heteroplasmy levels into wild-type (WT; 0 – 0.5%), low (0.5% – 30%), medium (30% – 65%), and high (65% – 100%) heteroplasmy, indicated by dashed lines. **(C)** Heatmap showing average mitochondrial RNA expression (z-score) across heteroplasmy categories (Zero, Low, Medium, and High). Genes deleted in mtDNA^ΔScaI^ are highlighted in red, whereas partially deleted genes are highlighted in pink. **(D)** UMAP of 25,509 mtDNA^ΔScaI^ cells in three media conditions. Cells are colored by growth conditions. UMAP was computed using mitochondrial-encoded genes (13) as features. **(E)** UMAP of 25,509 mtDNA^ΔScaI^ expressing cells color-coded by heteroplasmy levels. UMAP was computed using using mitochondrial-encoded genes as features. (**F)** Scatterplots of average log-scale mitochondrial RNA as a function of heteroplasmy in cells with the indicated treatments. Each dot represents an average of 20 cells ranked by heteroplasmy levels (see Methods for a detailed description of cell binning).

UMAP visualization, constructed from the top 4500 highly variable genes, separated cells cultured in galactose distinctly from those cultured in glucose with or without pyruvate (Figure S6C). However, when the analysis was restricted to mtRNA genes only, this separation became less pronounced (Figure 3D). Additionally, when we overlaid heteroplasmy data onto levels of all mtRNA, we observed that cells with similar heteroplasmy levels exhibited comparable mtRNA abundances (Figure 3E). This observation prompted further investigation of the relationship between heteroplasmy and the abundance of different mtRNAs. To prevent confounding effects from variations in mtDNA copy numbers, we developed an algorithm to estimate mtDNA content at the single-cell level. This allowed us to normalize RNA counts across all genes relative to mtDNA content (Figure S6D). We then superimposed normalized mtRNA levels and heteroplasmy scores at single-cell resolution (Figure 3F). The analysis revealed a gradual decline in mitochondrial transcripts from genes within the deleted region as the heteroplasmy levels rose. Genes outside the deleted region displayed less fluctuation at moderate heteroplasmies. However, beyond the critical heteroplasmy threshold, mitochondrial genes exhibited a sharp decrease in transcript levels, with the notable exceptions of ND1 and ND2, which accumulated significantly at high heteroplasmy levels (Figure 3F and Figure S6E). The unexpected late-stage enrichment of ND1 and ND2 was not observed in galactose conditions due to cell death at high heteroplasmy (Figure 3F). In contrast, the accumulation of mtDNA deletions did not significantly impact the expression of nuclear-encoded mitochondrial genes, confirming that the two transcriptional networks are independent (Figure S6F).

### Two distinct nuclear responses to increasing mtDNA^ΔScaI^ heteroplasmy

Next, we investigated the nuclear-encoded transcriptome in the DOGMAseq data to probe the nuclear transcription response to increasing mtDNA deletion by comparing gene expression in low, medium, and high heteroplasmy to wild-type cells (zero heteroplasmy) when cultured in different media. Differential gene expression analysis of each heteroplasmy group against wild-type revealed distinct gene expression changes as a function of heteroplasmy load and available metabolites (Figure 4A and Table S2). In addition to the anticipated transcriptional response at high heteroplasmy, we noted substantial gene deregulation in cells with medium mtDNA^ΔScaI^ burden (Figure 4A and S7A-B and Table S2). This observation suggests that cellular sensing and transcriptional rewiring occur even when cells accumulate mtDNA^ΔScaI^ below the critical threshold of heteroplasmy. Pathway analysis of top differentially expressed genes highlighted different biological pathways deregulated at various heteroplasmy levels, including Ca^2+^ sensing and cholesterol biosynthesis. Other pathways, such as extracellular matrix organization, were exclusive to a specific heteroplasmy level (Figure 4B and Table S3).

**Figure 4.**
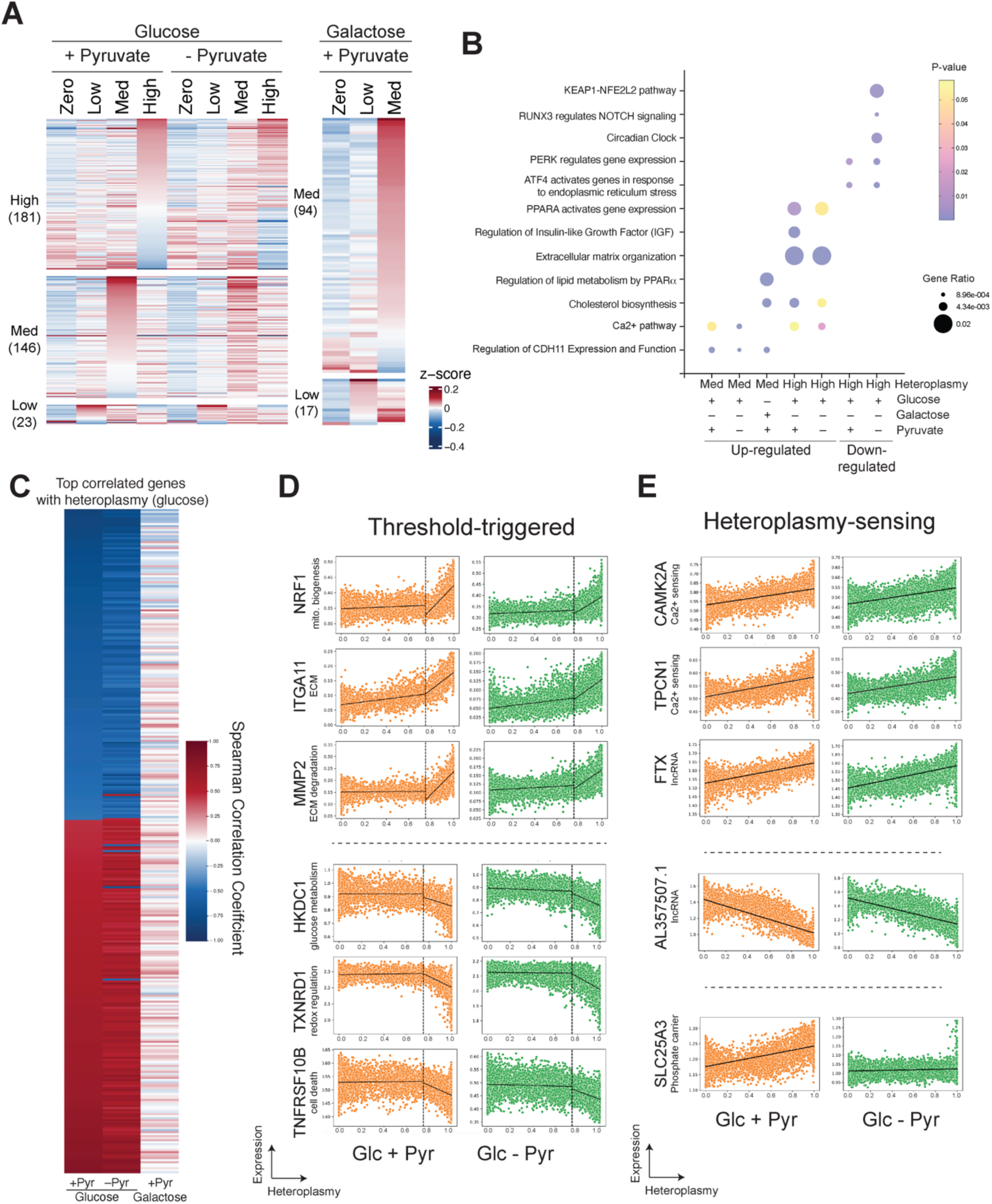
Evidence for two distinct nuclear responses to increasing mtDNA^ΔScaI^ heteroplasmy. **(A)** Heatmap depicting the average expression levels of differentially expressed nuclear-encoded genes in Low, Med, and High heteroplasmy cells compared to Zero cells with the indicated culture conditions. Colors indicate z-scores across heteroplasmy categories (Genes in Table S2). **(B)** Dotplot of selected biological pathways up or down-regulated across medium and high heteroplasmy categories. See Table S3 for a list of the pathways for each condition and treatment. Analysis done using www.reactome.org. **(C)** Spearman correlation heatmap of top genes correlated with heteroplasmy. Genes displayed with Spearman scores > 0.45 or < −0.45 in Glucose + Pyruvate and Glucose -Pyruvate. **(D)** Scatterplot of highlighted genes exhibiting a threshold-driven change in gene expression in response to high heteroplasmy. Plots display MAGIC imputed expression in single cells. **(E)** Scatterplot of highlighted genes that exhibit a linear increase or decrease in gene expression in response to increasing levels of mtDNA^ΔScaI^ per cell. Plots show MAGIC imputed expression in single cells. See the Methods section for 3-step filtering criteria to identify genes in each category (**E&F**).

The transcriptional response observed at deletion levels below the heteroplasmy threshold led us to explore gene expression across a continuous spectrum of heteroplasmy. To achieve this, we analyzed scRNA-seq gene expression as a function of heteroplasmy at the single-cell level, focusing on glucose-rich media. To better classify gene expression patterns relative to heteroplasmy, we implemented a three-step gene filtering process based on Spearman correlation and the slope at defined heterplasmy (See Methods section for details). Our analysis revealed two distinct nuclear response patterns to mtDNA^ΔScaI^ levels (Figure 4C-E and Table S4). First, we observed a core set of genes that became deregulated once cells accumulated the critical threshold of >75% mtDNA deletions. Upregulated genes in this category include Nuclear Respiratory Factor 1 (NRF1), a transcription factor that activates several metabolic genes^42,43^, as well as integrin alpha 11 (ITGA11), the transmembrane protein CSMD1, and matrix metalloproteinase-2 (MMP2) (Figure 4D and S7C). Downregulated genes at high heteroplasmy included thioredoxin reductase 1 (TXNRD1) and Hexokinase domain containing 1 (HKDC1) (Figure 4D and S7C and Table S4).

Importantly, our single-cell multiome experiments uncovered an independent gene network that correlated steadily with increasing heteroplasmy. Genes showing gradual increase or decrease as a function of increased mtDNA^ΔScaI^ included ones linked to intracellular calcium sensing and regulation, such as calcium/calmodulin-dependent protein kinase II alpha (CAMK2a) – a multifunctional serine/threonine-specific protein kinase required for normalizing intracellular Ca^2+^ homeostasis, CAMKMT, and TPCN1 (Figure 4C and E and S7D). Specific heteroplasmy-sensing genes, such as the mitochondrial phosphate carrier protein SLC25A3, showed no response in conditions lacking pyruvate (Figure 4C and E). Last, our analysis identified several long non-coding RNAs deregulated in response to mtDNA^ΔScaI^ accumulation, including FTX, AL357507.1, and NEAT1, a structural component of nuclear paraspeckle (Figure 4C-E and Figure S7C-D). Altogether, our analysis of DOGMAseq data provided unparalleled insight into the nuclear response to increasing mtDNA deletion levels, uncovering both a threshold-triggered response and a gradual heteroplasmy-sensing gene network.

### Applying mito-EJ to engineer diverse mtDNA deletions across different cell types

Having successfully demonstrated that mito-EJ combined with a mitochondria-targeted restriction enzyme is an effective strategy for generating single mtDNA deletions in epithelial cells, we further explored the robustness of this method and its applicability in engineering other disease-causing mtDNA deletions across various cell types. First, we sought to engineer smaller mtDNA deletions in epithelial cells. For this purpose, we expressed mito-ApaLI, which targets a single mtDNA cleavage site, along with mito-EJ components in *MGME1^-/-^*ARPE-19 cells. ATAC-seq analysis revealed an enrichment of mtDNA molecules with a small deletion (Figure 5A and S8A and Table S1). We also detected small insertions likely introduced by the polymerase activity of LigD disrupting the restriction site (Figure 5B and Table S1). Next, we leveraged mito-EJ to reconstitute the common deletion, a widespread 4.9kb deletion causing several mitochondrial syndromes, and enriched in aging tissues^44,45^. To achieve this, we employed TALENs to introduce breaks flanking the 5’ and 3’ ends of the common deletion (Figure 5C). qPCR analysis revealed a significant accumulation of the 4.9kb deletion in cells expressing mito-EJ compared to control cells (Figure 5D and S8B-D). Finally, we tested the two-component system in generating mtDNA depletions in other cell types, specifically human induced pluripotent stem cells (iPSCs), which can be differentiated into disease-relevant cell types. We stably expressed mitochondrial T4 DNA ligase and Dox-inducible mito-ScaI in iPSCs(Figure S8E-F). Upon Dox induction of ScaI expression, we detected a significant accumulation of mtDNA^ΔScaI^, particularly when enriching for cells expressing the cleaving enzyme (Figure 5E-F and S8F).

**Figure 5.**
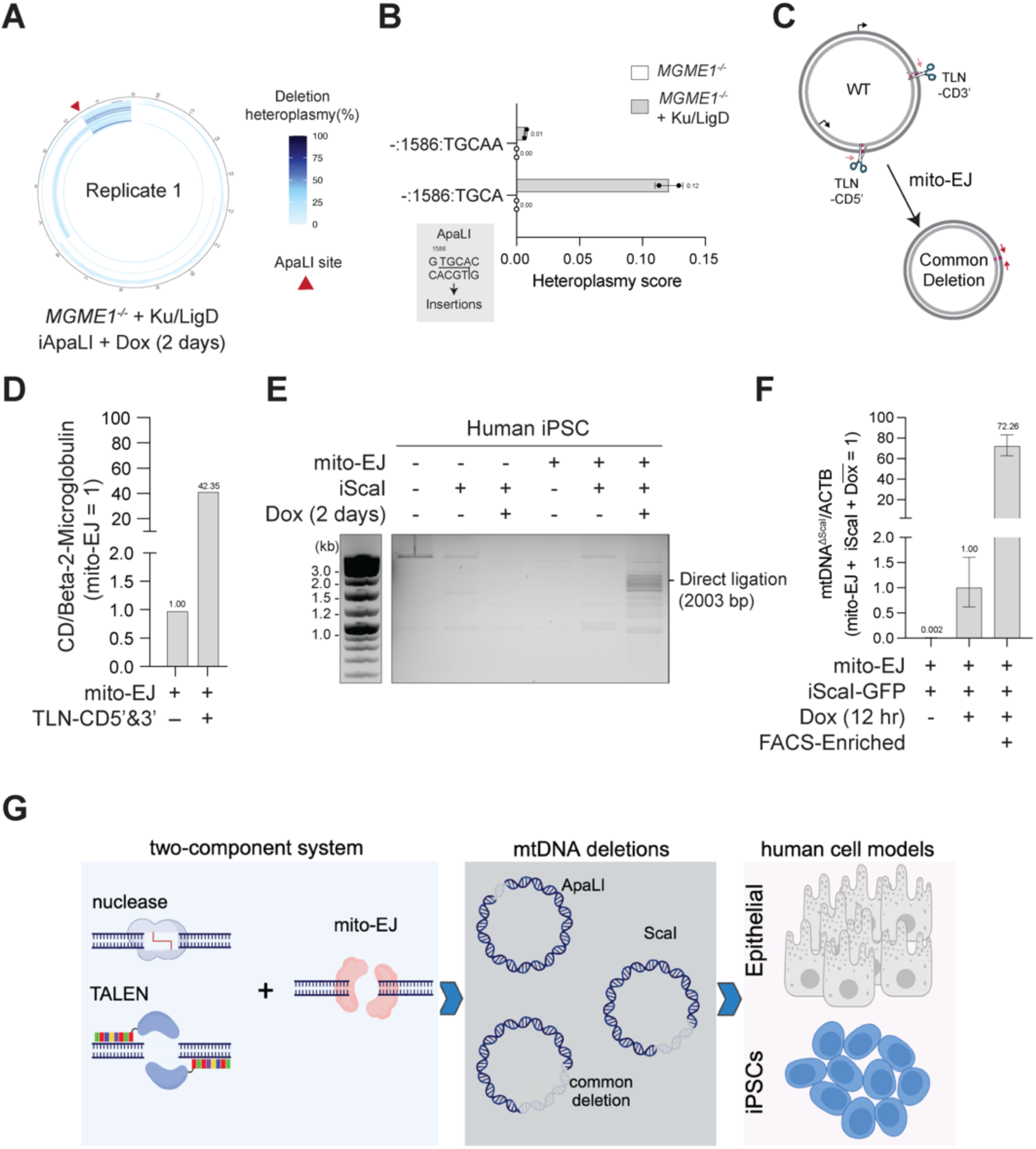
Applying mito-EJ to engineer diverse mtDNA deletions in various cell types. **(A)** Circular plots showing deletions identified based on ATAC-seq in *MGME1^-/-^* cells expressing mito-EJ (Ku & LigD) induced for mito-ApaLI (dox-inducible promoter). Cells were treated with Dox for 24 hours and recovered for four days. Table S1 depicts all deletions detected by ATAC-seq. **(B)** A graph highlighting heteroplasmy of small insertions was detected at the ApaLI site in *MGME1^-/-^* ARPE-19 cells with the indicated treatment. Bar plots are means ± s.d. from n = 2 replicates. Inserts are listed in Table S1 **(C)** Schematic of the 4977-bp deletion, named “common deletion (CD)”, engineered using mito-TALENs (TLN-CD3’ and TLN-CD5’) that targeting the 3’ and 5’ ends in cells expressing mito-EJ. Genotyping PCR primers are in red. **(D)** qPCR quantification of the level of common deletion (CD) in samples from (F), normalized to B2M locus in the nuclear DNA. The value of the mito-TALENs transfected sample is normalized to the untransfected sample. **(E)** PCR to detect the formation of mtDNA^ΔScaI^ in MSK-SRF001 human induced pluripotent stem cell (iPSC) expressing mito-T4 DNA ligase as the mito-EJ (T4 Ligase) system and Dox-inducible ScaI and treated with Dox treatment for two days. Highlighted is the PCR product from upon ligation of the two most distal ScaI sites. **(F)** qPCR quantification of mtDNA^ΔScaI^ levels in iPSCs expressing mito-EJ and Dox-inducible ScaI. Cells were treated with Dox for 12 hours and harvested on Day 4. The enriched sample was sorted based on ScaI-T2A-GFP expression. mtDNA^ΔScaI^ was normalized to ACTB. Normalized values are means ± s.d. from n = 3 technical replicates. **(G)** Schematic for applying the two-component systems of mito-EJ and programable nucleases to engineer specific mtDNA deletions in various cell types.

## DISCUSSION

### Implications and applications of the mtDNA deletion-generating tool

Since the first disease-causing mtDNA deletion was discovered in 1988, over 150 mtDNA deletions have been found in mitochondrial disorders and age-related diseases ^45^. Investigating the pathogenicity of large-scale mtDNA deletions and identifying treatment strategies has been challenging due to mosaicism, cell-type-specific manifestations of deletions, and, most significantly, the inability to engineer deletions in human cells. The successful implementation of the two-component system – mito-EJ and mito-nucleases – in human cells provides a powerful method to model pathogenic mtDNA deletions at defined heteroplasmy levels and in the desired cell types (Figure 5G). Inspired by findings based on the fly studies, where T4 ligase effectively prevented the degradation of cleaved mtDNA and facilitated deletion formation^13^, we reconstituted two prokaryotic end-joining systems within human mitochondria. We demonstrated that either Ku&LigD or T4 Ligase can overcome the organelle’s intrinsic lack of DSB repair, leading to the formation of mtDNA deletions in response to programmable nucleases. As a proof-of-concept, we applied this method to create a ∼3.5 kb mtDNA deletion and leveraged its cellular heterogeneity to isolate clonal cell lines spanning the entire range of heteroplasmy. These cells offer significant advantages over cybrids^18^, which exhibit a mismatch between nuclear and mitochondrial genomes, and iPSCs typically carry a single heteroplasmy burden dictated by the cell of origin^20^.

Using the two-component system, it is now feasible to model disease-relevant mtDNA deletions at defined heteroplasmy in various disease-related tissues. This is the first and necessary step toward deciphering the tissue-specific biochemical and pathophysiological impact of the deletion. Furthermore, by reconstituting the deletion in embryonic stem cells, one can compare how mtDNA mutations influence their differentiation into different lineages. In addition to generating single deletions, the mito-EJ system allows for the simultaneous induction of multiple deletions by combining different nucleases. This enables the investigation of the competition between various deletions, including ones lakcing individual single mitochondrial genes, tRNAs, or large-scale genomic regions. Creating a panel of epithelial cells with a full spectrum of heteroplasmy for a large-scale deletion is a critical tool for probing the metabolic and cellular signaling and transcriptional rewiring at different heteroplasmy levels. Moreover, this panel of cells provides a cellular model for early-stage drug testing and the development of treatments for mitochondrial disorders like progressive external ophthalmoplegia (PEO), Kearns-Sayre syndrome (KSS), Pearson marrow syndrome, and conditions associated with accelerated aging.

### mtDNA deletions – the threshold effect

Investigating genotype-phenotype relationships in clinical cohorts supported the existence of a critical threshold of mutant mtDNA, beyond which wild-type mitochondrial genomes can no longer buffer their pathogenic effects^46–48^. The threshold varies according to the mutation type and impacted tissue type. For example, while mutations in mtDNA can clonally expand and potentially reach homoplasmy, deletions are typically heteroplasmic^49^. Furthermore, tissues with high energy demand, including muscle and brain, are most commonly affected by mtDNA deletions ^50^. Nevertheless, what is the maximal burden of pathogenic mtDNA deletions, and how does the biochemical threshold translate into cellular defects and manifest clinically? Here, we began to address this gap in our knowledge by characterizing epithelial cells that carry the entire spectrum of mtDNA deletion heteroplasmy.

We identified a threshold of approximately 75% mtDNA deletions, beyond which retinal pigmental epithelial cells display severe metabolic impairment and cellular dysfunction. Our findings show that cells surpassing this heteroplasmy threshold experience significant depletion of most OXPHOS components (Figure 2B). The depletion of respiratory subunits encoded by genes retained in mtDNA^ΔScaI^ is likely due to defective mitochondrial translation caused by the loss of three tRNAs in the deleted region. We observed a concomitant loss of nuclear-encoded OXPHOS proteins, which appeared independent of transcription and not caused by impaired mitochondrial protein import, especially since TFAM and VDAC1 were not degraded. Given that mitochondrial and cytoplasmic translation programs are not coupled^51^, depletion of nuclear-encoded OXPHOS subunits near homoplasmy is likely due to increased protein degradation upon failure to integrate into a respiratory supercomplex. Coupled with severe defects in OXPHOS, cells with >75% heteroplasmy exhibited disruption in redox balance and impaired cell growth upon glucose withdrawal (Figure 2 and S4). Ultimately, while our study offered in-depth insights into mtDNA deletion outcomes in epithelial cells, employing the two-component system will be instrumental for comparing deletion thresholds and metabolic effects across different cell types, particularly post-mitotic cells.

In addition to the observed metabolic and cellular dysregulation, we discovered a significant upregulation of ND1 and ND2 in cells with high heteroplasmy. While an overall increase in mtRNA in cells containing mtDNA mutations has been previously reported ^52^, the selective upregulation of ND1 and ND2 genes has not been observed. mtDNA is transcribed into a long polycistronic RNA that is cleaved into individual components and processed co-transcriptionally. Thus, the regulation of ND1 and ND2 is likely to be post-transcriptionally. In support of this notion, quantitative analysis showed that the abundance of mitochondrial transcripts is primarily dictated by the stability of mitochondrial mRNA^53^. A key player in this process is LRPPRC, a protein associated with the metabolic disorder Leigh syndrome, which is critical for the stability, translation, and folding of mtRNA^53–55^. It is plausible that the selective stabilization of ND1 and ND2 results from altered binding to LRPPRC in response to mitochondrial dysfunction. Future studies must investigate the causes and consequences of ND1 and ND2 accumulation. It is imperative to determine whether this upregulation acts as a compensatory mechanism for the loss of oxidative phosphorylation (OXPHOS) or indicates a potential non-canonical function activated by the accumulation of deletions.

Interestingly, unlike in *C. elegans*, where the emergence of mtDNA deletions leads to their self-propagation by increasing mtDNA replication, the mtDNA^ΔScaI^ heteroplasmy in ARPE19 clones remained stable in glucose-containing media (Figure S3E). Instead, metabolic stress selectively eliminated mutant mtDNA (Figure S3E). These results are consistent with reduced cell fitness triggering a cell-level selection similar to data obtained from cells expressing a nonsense mutation in a complex I subunit ^56^. Alternatively, we cannot rule out the possibility that organelle and nucleoid-based selection might also operate to eliminate mtDNA deletions in somatic cells under metabolic stress. Using mito-EJ to reconstitute mtDNA deletions in different cell lineages, we can now start to interrogate the underlying basis of mtDNA deletion propagation in somatic cells. Importantly, the method will help uncover tissue-specific dynamics of mtDNA deletions in cells with high energy demands, such as neurons and muscle cells, which primarily manifest mitochondrial pathologies due to mtDNA deletions.

### Two-tier nuclear sensing of mtDNA deletions

Analysis of cybrids comprising a point mutation in mtDNA highlighted abrupt changes in transcriptional activity at critical heteroplasmy levels ^52^, leading to the assumption that a nuclear response is triggered upon severe mitochondrial dysfunction. Investigating mouse models of mtDNA deletions revealed a disease stage activation of the integrated stress response (ISRmt) controlled by mTORC1 and involving FGF21 and other metabolic pathways that drive tissue remodeling^57,58^. A thorough and stepwise recording of the nuclear response to the continuous spectrum of deletion heteroplasmy, though currently lacking, is essential to reveal the cellular effects of mtDNA deletions. Our single-cell multiomic approach provided insight into gene expression programs as a function of heteroplasmy and in different culture conditions. Data obtained in galactose-containing media has limitations, particularly since high levels of mtDNA^ΔScaI^ triggered cell death. Instead, findings based on glucose-rich media revealed distinct gene networks deregulated at various heteroplasmy levels. Beyond the anticipated threshold-triggered transcriptional response, we observed a gene expression profile consistent with sensing of heteroplasmy at levels below the critical threshold to maintain a viable level of oxidative phosphorylation (Figure 4C and E). Further supporting the gradual sensing of mtDNA deletions, Seahorse analysis of clones with increasing heteroplasmy shows a graded impairment in oxygen and proton flux (Figure 2C). These observations raise the question of how cells detect mtDNA deletions. The constituent upregulation of factors linked to the regulation of calcium levels and signaling, including CAMK2a, CAMKMT, AHNAK2, and TPCN1(Figure 4D and E and Figure S7D)^59^, hints at deregulated Ca²⁺ signaling – previously linked to mitochondrial respiratory chain diseases – as a potential sensor of mtDNA deletions^60–62^. Our single-cell level characterization of mtDNA deletion-triggered gene networks establishes a foundation for future studies to interrogate the sensing mechanisms within clinically relevant tissues. This groundwork also paves the way to explore the effects of manipulating these pathways on disease manifestations and potential therapeutic interventions.

### Limitations of the study

In this study, the dynamics of mtDNA deletion, pathological threshold, and transcriptional response were characterized using an *in vitro* culture system comprising a retinal pigment epithelial cell line. The advantage of this normal diploid cell line is that, when cultured in glucose, it relies on glycolysis, allowing the accumulation of mtDNA deletion to near homoplasmy. When the system is shifted to stress conditions, the effect of the deletion is unmasked. Nevertheless, the threshold and response will likely vary in different cell types, including postmitotic cells and *in vivo*. In that respect, implementing the two-component genetic tool to induce large-scale deletions in cells with high energy demand or embryonic stem cells that can then be differentiated into different lineages will help unravel the impact of the deletion in a range of disease-relevant tissues.

## ACKNOWLEDGEMENTS

We thank Caleb Lareau for advice on the computational analysis of DOGMAseq and Carlos Moraes for providing the mito-ApaLI and mito-ScaI plasmids. We thank members of the Sfeir lab for technical and intellectual support and feedback on the manuscript. This work was supported by NIH/NIA (R01AG085782) and Pew Charitable Trust grants, The G. Harold & Leila Y. Mathers Charitable Foundation, and the Edward Mallinckrodt Jr. Foundation Award (A.S.). M.T. is supported by the NIH/NIGMS (R35GM147191). This work is supported by P30CA008748 (MSKCC). T.K. is the Timmerman Traverse Fellow of the Damon Runyon Cancer Research Foundation (DRG-2521-24). Work in the C.B.T laboratory is supported by the NIH/NCI (R35CA283988). The Single Cell Analytics and Innovation Lab performed all Single-Cell Genomics: M.L, I.M, R.S, M.T., R.S, and R.C. were supported by the Gerry Metastasis and Tumor Ecosystems Center. D.P. is a Howard Hughes Medical Institute Investigator.

## AUTHOR CONTRIBUTION

Conceptualization: A.S. and Y.F.

Methodology: A.S., Y.F., M.L.

Investigation: Y.F. with help from T.K., R.C., T.L., K.W.R., M.T., T.L. E.D., I.M., D.B., M.T.

Single-cell genomics I.G., M.T., R.C.

Data Analysis: M.L., R.S., R.S., R.C., M.K., Y.F., D.P.

Visualization: Y.F., T.K., T.L.

Funding acquisition: A.S., C.B.T., D.P, E.R., M.T.

Project administration: A.S. Supervision: A.S.

Writing – original draft: A.S., with help from Y.F. and M.L.

Writing – review & editing: all authors

## Supplemental Material

### Supplemental Figures

**Figure S1.**
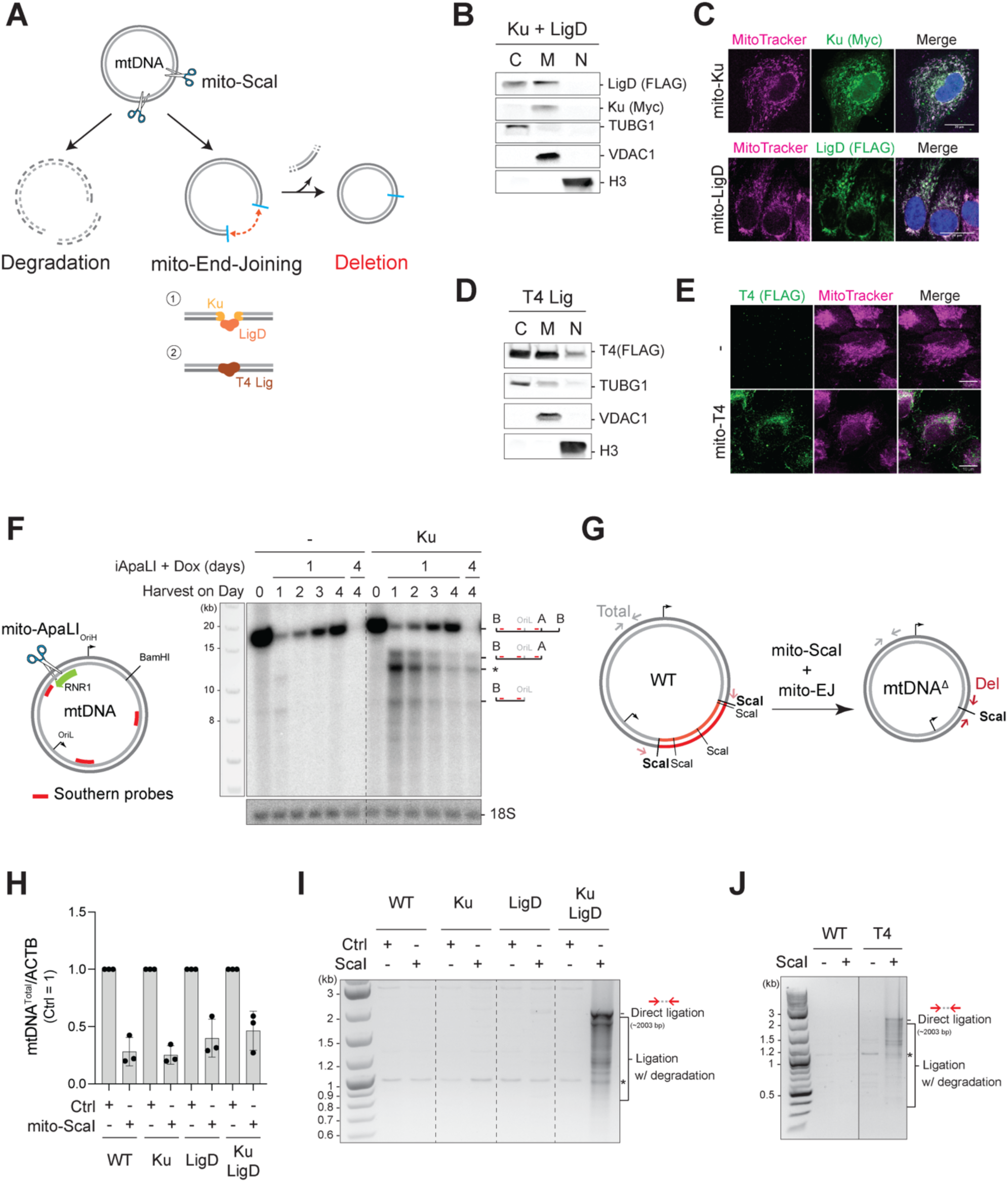
Reconstitution of end-joining (EJ) in the mitochondria. **(A)** Schematic of engineering mtDNA deletion via end joining. Cleaved mtDNA is primarily subject to degradation. Introducing DNA end joining systems, such as 1) Ku and DNA ligase LigD, and 2) T4 DNA ligase, into mitochondria could ligate DNA ends, leading to the loss of intervening sequence and the accumulation of mtDNA deletions. **(B)** Western blot analysis for expression and localization of Ku and LigD following subcellular fractionation into cytoplasmic (C), mitochondrial (M), and nuclear (N) fractions. **(C)** Immunofluorescence for Ku and LigD in ARPE-19 cells. Mitochondria are marked with Mitotracker. **(D)** Western blot analysis for expression and localization of T4 ligase following subcellular fractionation. **(E)** Immunofluorescence for T4 ligase. Mitochondria are marked with Mitotracker. **(F)** Left, schematic depicting the mito-ApaLI cut site on mtDNA. BamHI was used to linearize DNA for southern blot analysis. Probes to detect mtDNA are highlighted in red. Right, southern blot analysis of linear mtDNA fragments in cells expressing mito-Ku and subjected to induction of mito-ApaLI on Day 0. A, ApaLI. B, BamHI. OriL, replication origin of the light strand. **(G)** Schematic depicts primers that distinguish deletions formed following mito-ScaI cleavage and mito-EJ from wild-type (WT) mtDNA and primers for detecting total mtDNA. **(H)** qPCR analysis of relative mtDNA copy number in cells expressing indicated components of mito-EJ and transfected with mito-ScaI or control plasmid, as in Figure 1B (left). Plotted values are mean ± s.d. from *n* = 3 technical replicates. **(I)** PCR to detect ligation activity of Ku and LigD expressed and localized to mitochondria upon transfection of mito-ScaI or a control plasmid for four days. Direct ligation indicates the approximate size of PCR for ligation between the most distal ScaI sites. Ligation with degradation indicates larger deletions due to the degradation of DNA ends. **(J)** PCR is used to detect the ligation activity of T4 DNA Ligase, which is expressed and localized to mitochondria upon transfection of mito-ScaI or a control plasmid for four days.

**Figure S2.**
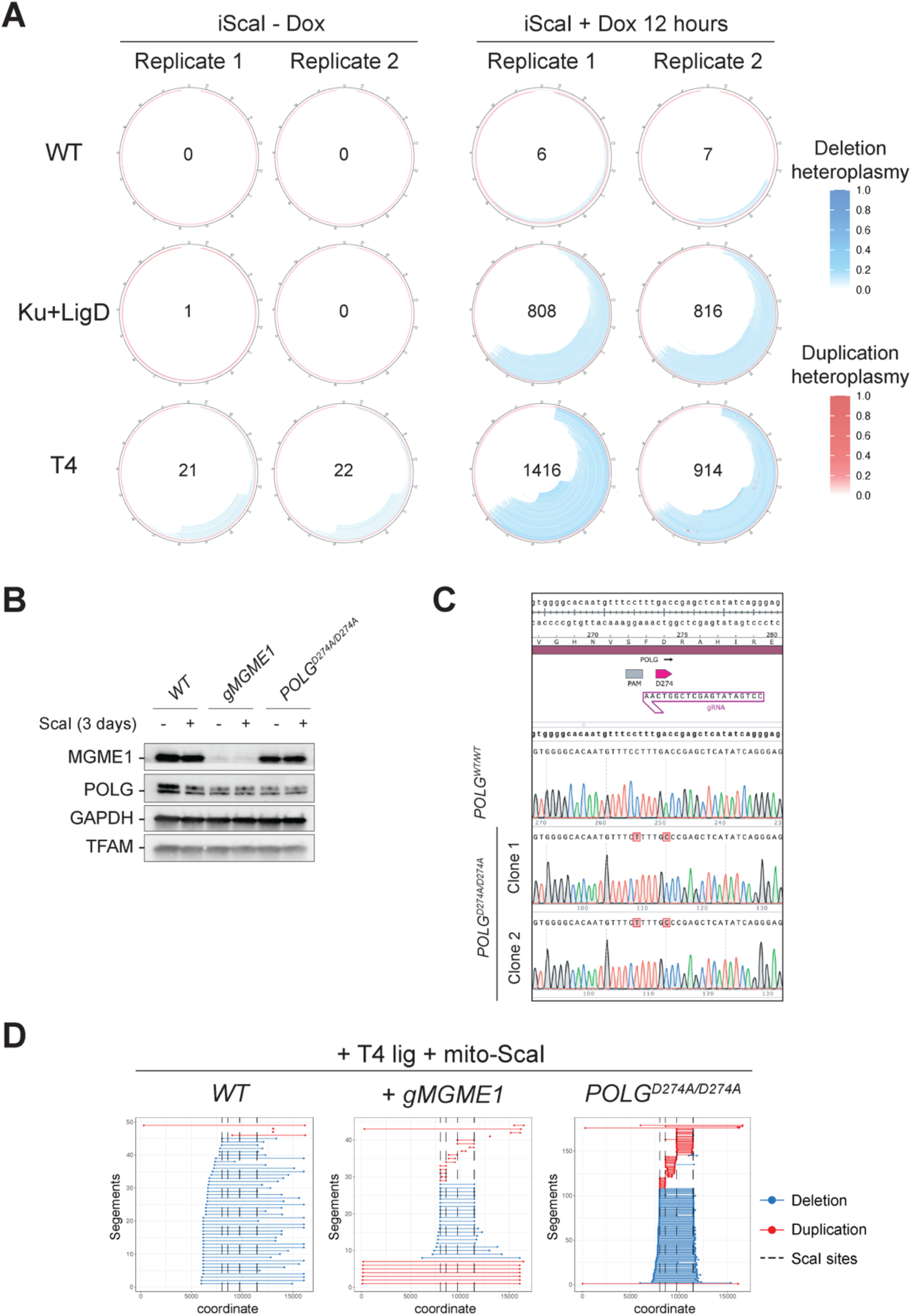
mtDNA deletion in cells treated with mito-ScaI and mito-EJ. **(A)** Circular plots showing mtDNA deletion accumulation in two independent replicates in cells expressing Ku & LigD, T4, and control cells. ARPE-19 cells expressing mito-EJ and doxycycline (Dox) inducible mito-ScaI were treated with Dox for 12 hours and harvested on Day 5 for ATAC-seq. The number of unique deletion types detected is highlighted in the center of each circular plot. **(B)** Western blot analysis for MGME1 in ARPE-19 cells treated with CRISPR/Cas9 to delete MGME1 (pooled cells) and POLG in clonally derived cells targeted with CRISPR/Cas9 to introduce p.D274A mutation in the exonuclease domain. GAPDH and TFAM serve as a control. **(C)** Sanger sequencing of three representative homozygous clones to confirm the POLG mutations at p.D274A and the PAM site. **(D)** Graphs displaying deleted (blue) and duplicated (red) mtDNA regions in cells with the indicated genotype and co-expressing of mito-ScaI and T4 ligase. Graph corresponds to data presented in Figure 1D.

**Figure S3.**
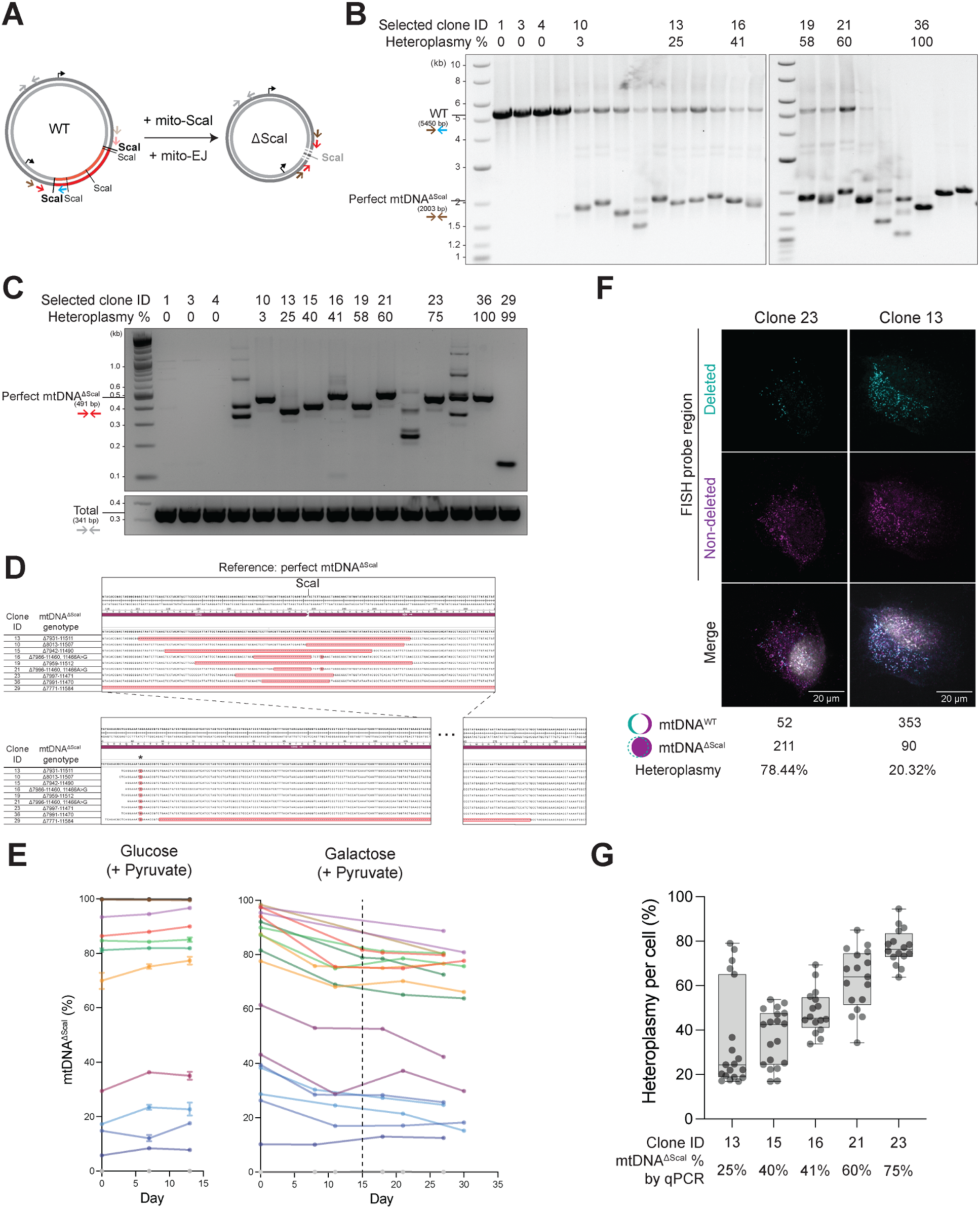
Isolation and characterization of mtDNA^ΔScaI^ clones. **(A)** Schematic of primers used to genotype wild-type and deleted mtDNA. **(B)** Agarose gel electrophoresis of genotyping PCR products using indicated primers to screen for clones carrying a single deletion. Three primers were used to amplify both wild-type and deleted mtDNA. Labeled clones were selected for in-depth studies. **(C)** Agarose gel electrophoresis of genotyping PCR products using indicated primers to screen for clones carrying a single deletion. Labeled clones were subsequently used. **(D)** The mtDNA^ΔScaI^ genotypes of selected clones were determined by Sanger sequencing and alignment to the perfect mtDNA^ΔScaI^ reference genome. Clone #29 carried a larger mtDNA deletion that is presented in split windows. * indicates a mutation preexisting in the parental cell line. **(E)** Heteroplasmy levels of mtDNA^ΔScaI^ in individual clones grown in glucose versus galactose medium for 13 or 30 days, respectively. Heteroplasmy was measured by qPCR. **(F)** Representive images of mtDNA FISH signals in individual cells and the corresponding quantification of wild-type and deleted mtDNA and heteroplasmy levels. **(G)** Quantification of single-cell heteroplasmy of mtDNA^ΔScaI^ in intermediate heteroplasmic clones using mtDNA FISH. Each dots represent individual cells, and 16-19 cells were analyzed per clone.

**Figure S4.**
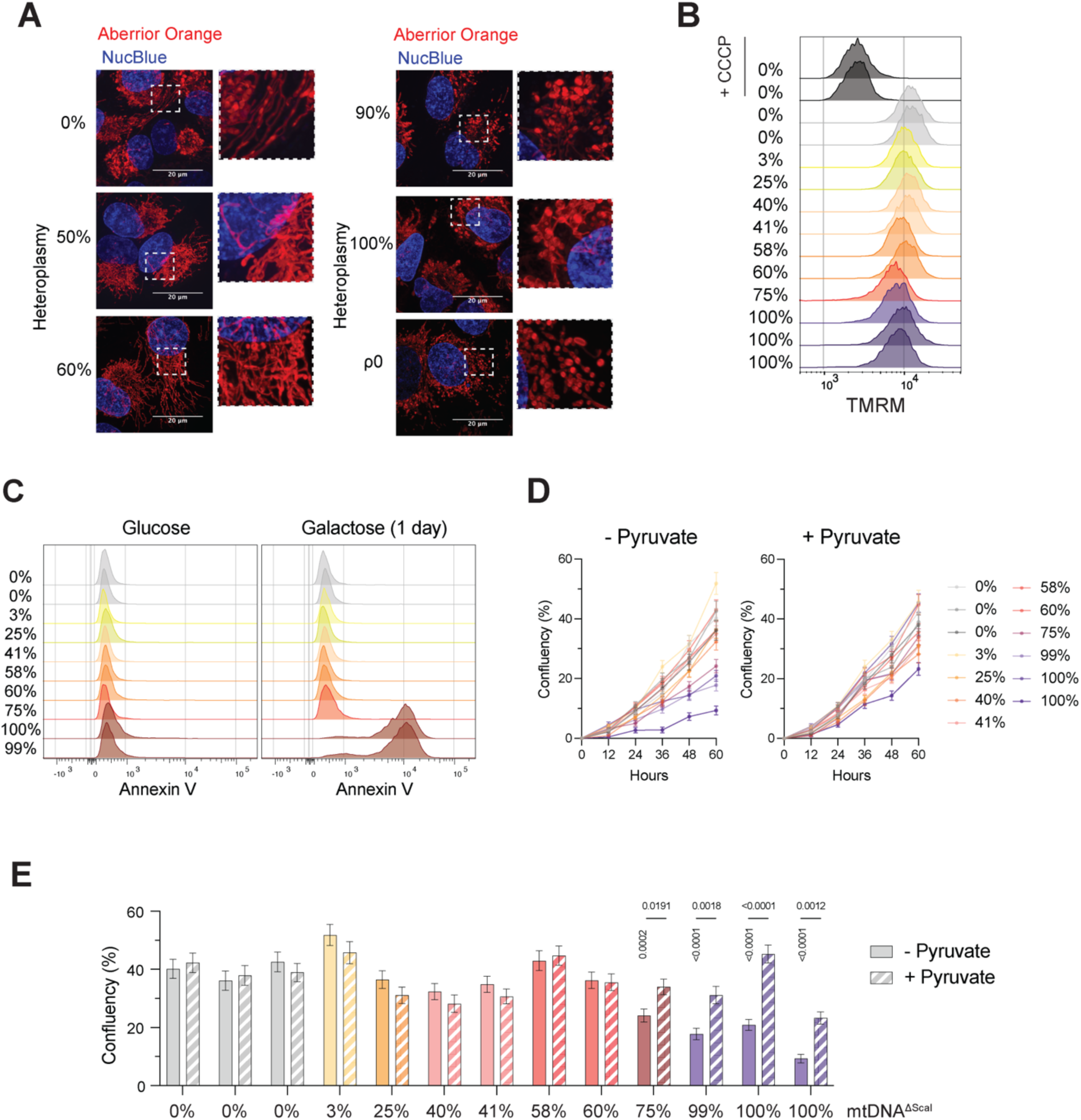
mtDNA deletion in cells treated with mito-ScaI and mito-EJ. **(A)** Representative SoRa spinning-disk confocal images of mitochondrial morphology in mtDNAΔScaI and ρ0 ARPE19 cells. Scale bars, 20 μm. Nuclei were labeled with NucBlue™ Live ReadyProbes™ (Hoechst 33342) and anterior LIVE ORANGE stained mitochondria. (**B)** Histogram of TMRM signals measured by flow cytometry to indicate mitochondrial membrane potential. CCCP treatment in 0% mtDNA^ΔScaI^ clones for 6 hours is a positive control. **(C)** Histogram of Annexin V signals measured by flow cytometry to indicate apoptotic status. **(D)** Growth curves of mtDNA^ΔScaI^ clones in glucose – pyruvate and glucose + pyruvate media for 60 hours, measured by cell confluency using Incucyte. Plotted values are means ± s.d. from *n* = 3 technical replicates. **(E)** Bar plot showing the cell confluency as in **(D)** at 60 hours. Vertical p-values are compared to the first 0% clone in glucose – pyruvate medium. Horizontal p-values compare glucose –pyruvate and glucose + pyruvate media within the same clones.

**Figure S5.**
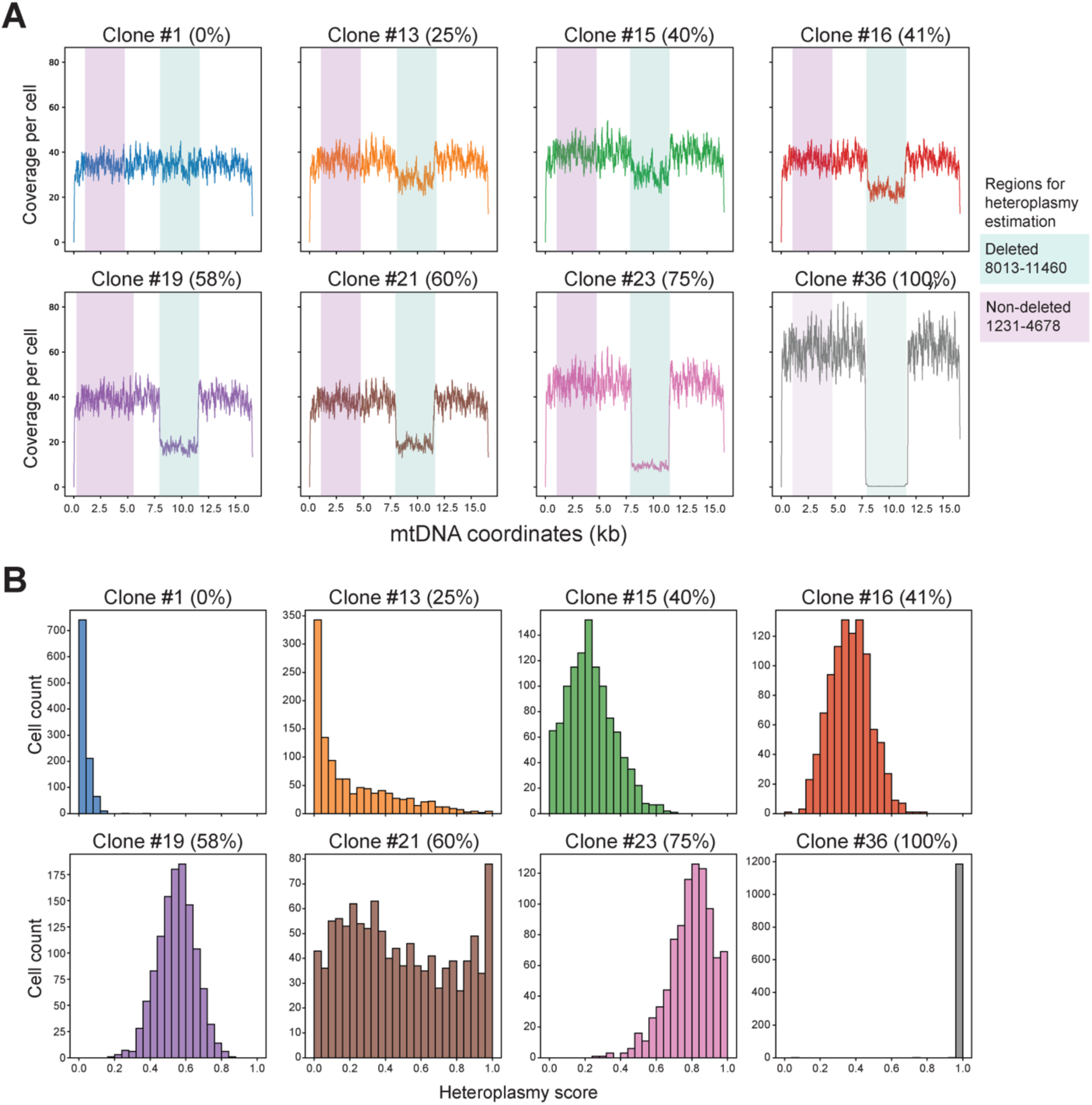
Eight clones of defined heteroplasmies were sequenced using DOGMAseq. **(A)** The average coverage per nucleotide per cell across the mitochondrial genome for each clone is depicted. We define coverage at a specific nucleotide as the number of distinct (unique start and end) DOMGA-seq ATAC reads that overlap it within a single cell. We report the average coverage per nucleotide for all cells per sample. High heteroplasmy clones have fewer reads (lower coverage) in the deleted region. **(B)** The cell distribution of estimated heteroplasmy for each clone, with the expected mean heteroplasmy measured by qPCR noted in parenthesis. Heteroplasmy is computed as one minus the ratio of deleted region to undeleted region (see Methods). Apart from Clone#21 (an outlier), the computed heteroplasmy score matched what was determined by q-PCR and FISH.

**Figure S6.**
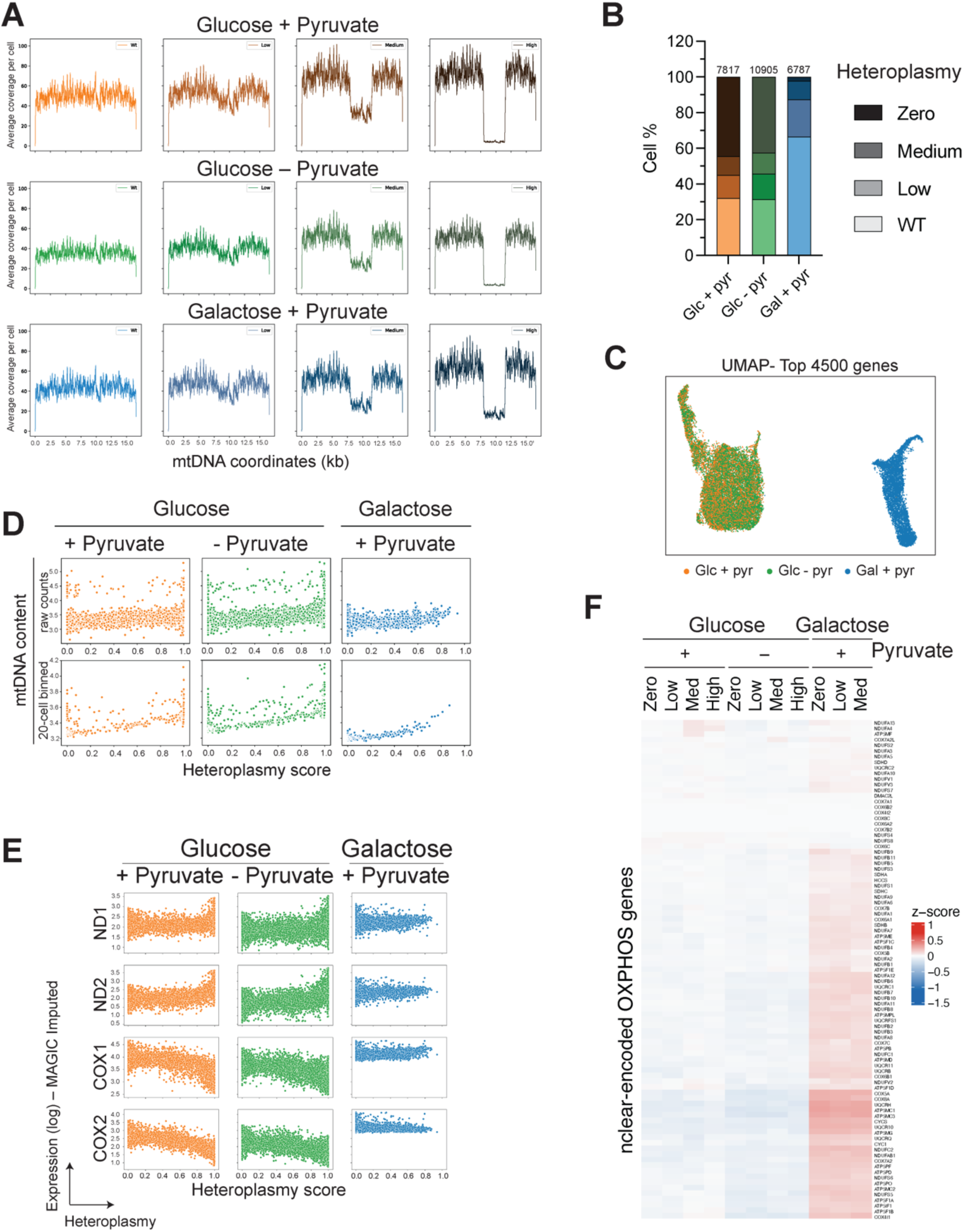
Overview of heteroplasmy distribution and copy number in Glucose + Pyruvate, Glucose – pyruvate, and Galactose + Pyruvate samples. **(A)** Average coverage per cell across the mitochondrial genome for each sample after binning cells into wild-type (defined as zero heteroplasmy) Low, Medium, and High heteroplasmy categories. **(B**) Bar graph depicting the percentage of cells in Zero, Low, Medium, and High heteroplasmy categories for each sample. **(C)** UMAP of 25,509 mtDNA^ΔScaI^ cells in three media conditions. Cells are colored by growth conditions. UMAP was computed using top 4500 genes. (**D**) Scatterplots of relative mtDNA content as a function of heteroplasmy score in samples with the indicated treatment. The top graphs are computed for cells, and the bottom plot shows mtDNA estimation after averaging every 20 cells. **(E)** Scatterplots of select mtRNA expression as a function of heteroplasmy in cells with the indicated treatments. Displayed in the MAGIC imputed expression *vs.* heteroplasmy levels corresponding to Figure 3F. **(F)** Heatmap illustrating the expression of nuclear-encoded mitochondrial-targeted proteins (z-score) for Zero, Low, Medium, and High heteroplasmy.

**Figure S7.**
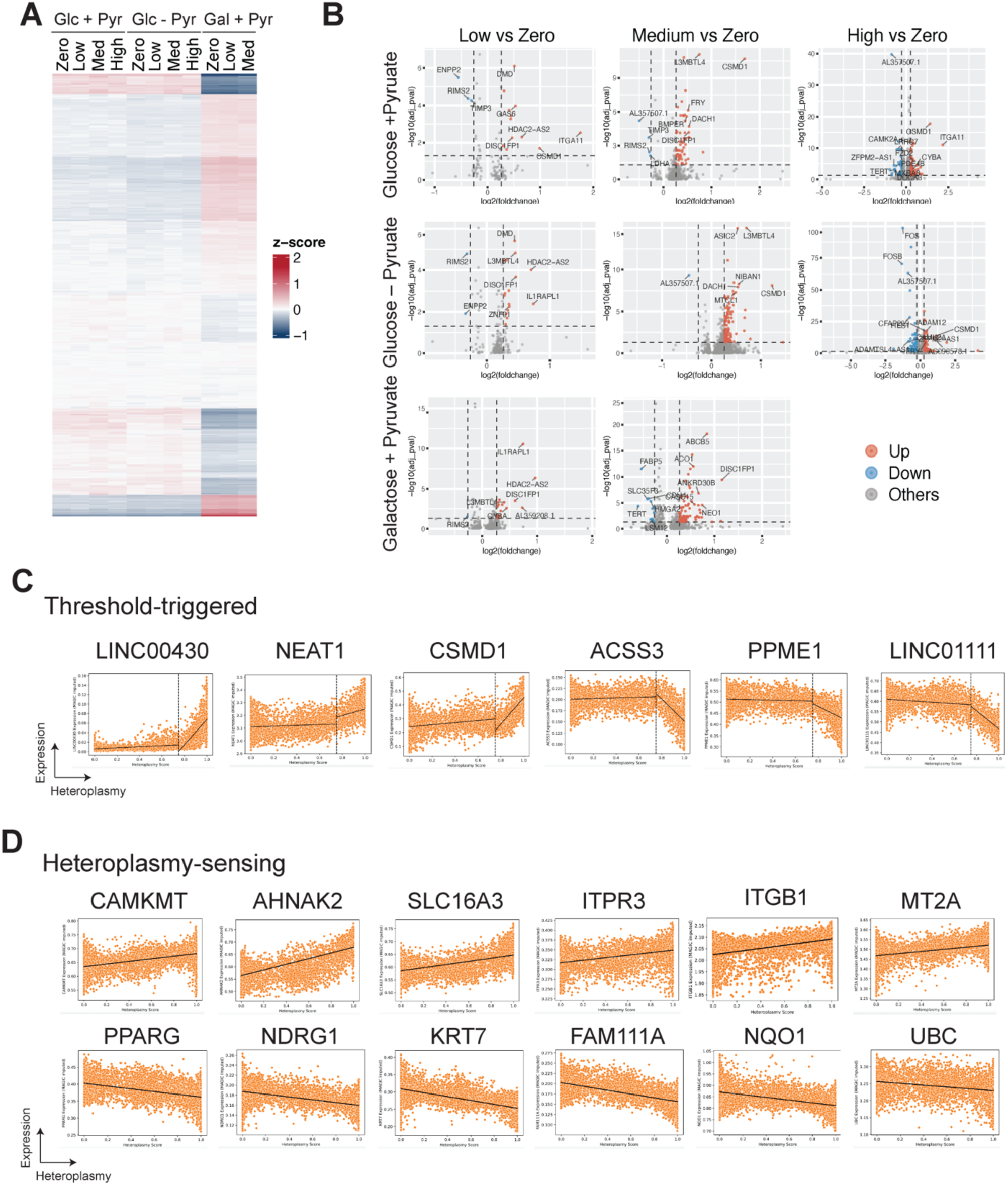
Differentially expressed genes *vs.* heteroplasmy levels in cells cultured under different conditions. **(A)** The heat map shows that the average expression of differentially expressed genes in cells is low, medium, or high heteroplasmy compared to wild-type in the indicated media condition. Colors indicate z-scores across all heteroplasmy categories. **(B)** Volcano plots of differentially expressed genes in Low, Medium, and High heteroplasmy compared to wild-type cells (zero heteroplasmy) using 1.2 fold change and 0.05 adjusted P-value as cut-offs. **(C-D)** Scatterplots of selected nuclear-encoded genes showing MAGIC imputed expression *vs.* heteroplasmy levels exhibiting thresholded-driven responses **(C)** and gradual deregulation upon sensing increasing heteroplasmy **(D)**. See Methods for filtering criteria.

**Figure S8.**
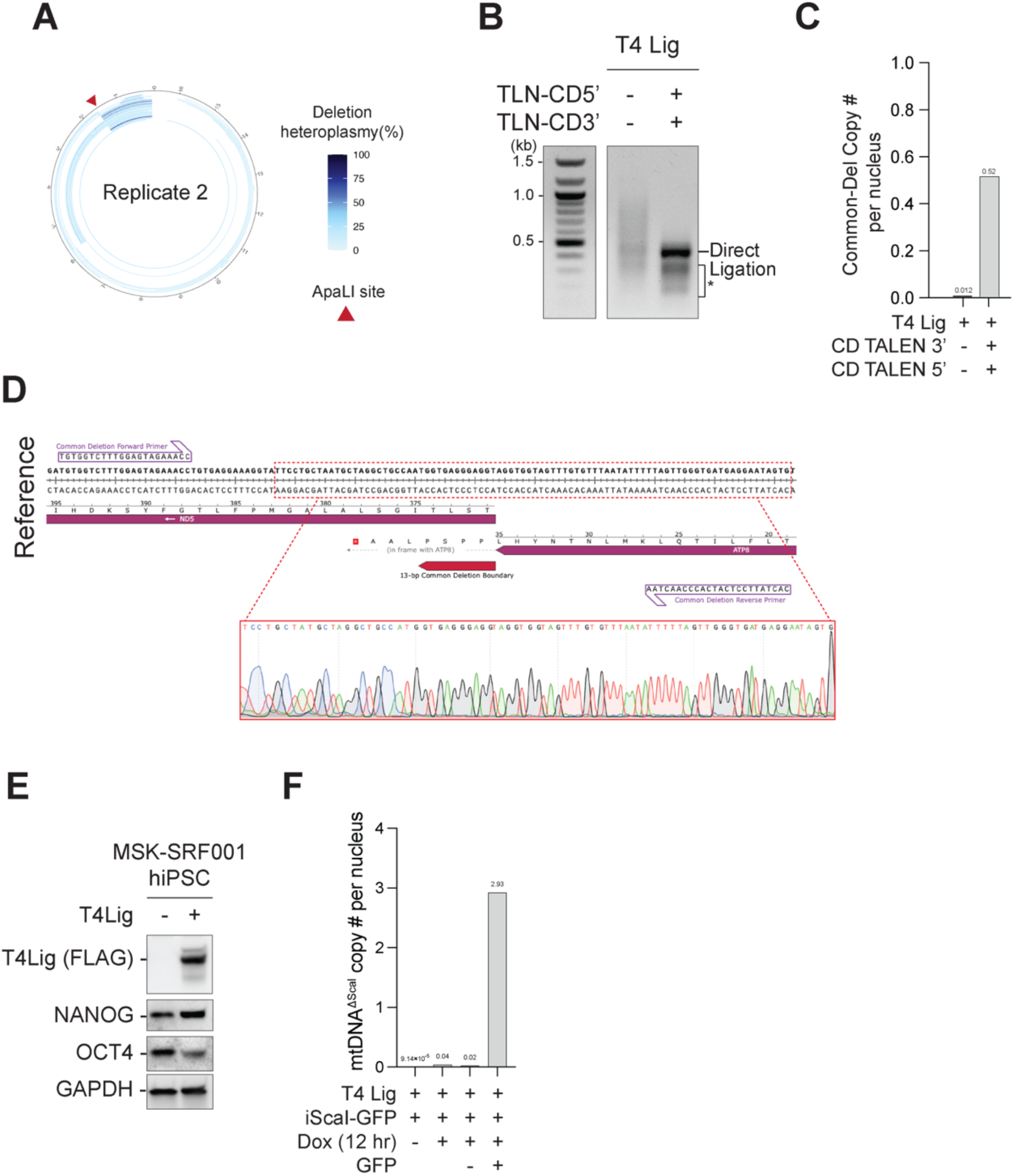
Applying mito-EJ to engineer different mtDNA deletions in various cell types. **(A)** Circular plots showing deletions identified based on ATAC-seq in *MGME1^-/-^* cells expressing mito-EJ (Ku & LigD) induced for mito-ApaLI (dox-inducible promoter). Cells were treated with Dox for 24 hours and recovered for four days. Replica #2 is shown (**B**) Agarose gel electrophoresis analysis of PCR products to detect the formation of the common deletion in ARPE-19 cells stably expressing mito-T4 DNA ligase and co-transfected with mito-TALENs (TLN-CD3’ and TLN-CD5’). Direct ligation indicates the size of PCR product from common deletion, and the smear below (*) indicates potential larger deletions due to degradation at the ligation junction. **(C)** Estimation of mtDNA Common Deletion copies per nucleus in samples as in Figure 5G. **(D)** Sanger sequencing of amplicons from genotyping PCR product in Figure 5F aligned to the sequence of Common Deletion. **(E)** Western blots analysis to detect mito-T4 DNA ligase and pluripotency markers NANOG and OCT4 in human iPSCs used in Figure 5A. **(F)** Estimation of mtDNA^ΔScaI^ copied per nucleus in samples used in Figure 5F.

**Table S1:** Deletions and insertions identified in bulk ATAC-seq and related to Figures 1 and 5.

**Table S2:** DEGs for heteroplasmy_vs_zero for all all_samples. Related to Figure 4A

**Table S3:** Enriched biological pathways for top deregulated nuclear genes (Heteroplasmic_vs_WT_ DEGs) based on Reactome analysis https://reactome.org/. Related to Figure 4B

**Table S4:** Spearman correlation and heteroplasmy sensing vs. threshold triggered genes. For tabs with Spearman correlation, the unfiltered tab lists all genes. filtered tab represent genes in Glucose + Pyruvate and Glucose – Pyruvate with spearman correlation > abs(0.45). Other tabs include nuclear genes identified as heteroplasmy-sensing and threshold-triggered according to the three-step filtering described in the methods section. Related to Figure 4C – E

**Table S5:** Oligonucleotides used in this study.

### Experimental Model and Study Participant Details

#### Cell culture procedures and treatment

Human retinal pigment epithelial cell line ARPE-19 cells (ATCC CRL-2302) were immortalized with hTERT. Before introducing dox-inducible transgenes, ARPE-19 cells and 293T cells (ATCC CRL-3216) were cultured in Dulbecco’s modified Eagle medium (DMEM, Corning, 10-013-CV) supplemented with 10% Bovine Calf Serum (GeminiBio), 2 mM L-glutamine (Gibco), 100 U/ml Penicillin-Streptomycin (Gibco), 0.1 mM MEM non-essential amino acids (Gibco), and 50 μg/ml uridine (Sigma). All ARPE-19 cell lines integrated with Dox-inducible mito-ScaI or mito-ApaLI were cultured in “Tet-free medium” which is Dulbecco’s modified Eagle medium (DMEM, Corning, 10-013-CV) supplemented with 10% tetracycline-free fetal bovine serum (FBS, TaKaRa 631368), 2 mM L-glutamine (Gibco), 100 U/ml Penicillin-Streptomycin (Gibco), 0.1 mM MEM non-essential amino acids (Gibco), and 50 μg/ml uridine (Sigma). In experiments where ARPE-19 cells were switched to different media, cells were seeded in Tet-free medium and on the next day, rinsed by PBS and switched to media based on DMEM without glucose (Gibco 11966025) and dialyzed fetal bovine serum (Gibco 26400044). Specifically, “glucose + pyruvate medium” is composed of DMEM without glucose (Gibco), supplemented with 10% dialyzed FBS (Gibco), 25 mM D-(+)-Glucose (Sigma G7021), 1 mM sodium pyruvate (Gibco 11360070), 2 mM L-glutamine (Gibco), 100 U/ml Penicillin-Streptomycin (Gibco), 0.1 mM MEM non-essential amino acids (Gibco), and 50 μg/ml uridine (Sigma); “glucose - pyruvate medium” is the same as glucose + pyruvate medium except without sodium pyruvate; “galactose medium” is the same as glucose + pyruvate medium except that 25 mM D-(+)-Glucose is replaced with 10 mM D-(+)-Galactose (Sigma G5388). MSK-SRF001 human induced pluripotent stem cells (iPSCs)^63,64^ were generously provided by the lab of Lorenz Studer (Memorial Sloan Kettering Cancer Center). iPSCs were maintained in Essential 8 medium (Gibco A1517001) in feeder-free conditions onto Matrigel-coated plates. Matrigel was purchased from MSKCC Stem Cell Research Facility and used following a published protocol^65^. iPSCs were passaged as clumps with 0.5 M EDTA in PBS every 4-5 days. All cells were grown in an incubator at 5% CO2 and 37°C. Cells expressing Dox-inducible mito-ScaI or mito-ApaLI were induced with 1 μg/ml doxycycline (Dox). Plasmids were transfected into ARPE-19 cells via a 4D-Nucleofector X unit (Lonza) using SF Nucleofection Buffer and the ER-100 program.

### METHOD DETAILS

#### Plasmids

mito-ScaI plasmid (pcDNA-mitoScaI) ^32^ was generously provided by the lab of Carlos T. Moraes (Miami School of Medicine). The doxycycline-inducible mito-ScaI plasmid (TLCV2-ScaI-T2A-EGFP-Puro) was cloned into the all-in-one Dox-inducible lentiviral backbone TLCV2 backbone (Addgene Plasmid 97360) with T2A-EGFP in-frame with mito-ScaI. Dox-inducible mito-ApaLI lentiviral plasmids (Addgene Plasmid 211462) were constructed in a previous study^26^. Genes encoding *Mycobacterium tuberculosis* Ku (Uniprot P9WKD9) and LigD (Uniprot P9WNV3) and enterobacteria phage T4 DNA ligase (Uniprot P00970) were codon-optimized for vertebrate, synthesized as gblocks (IDT), PCR amplified and cloned into various lentiviral plasmid backbones with addition of mitochondrial targeting signal of pyruvate dehydrogenase E1 subunit alpha 1 (PDHA1) at the N-terminus.

#### Cell line generation

ARPE-19 and MSK-SRF001 human iPSC cell lines stably expressing indicated transgenes were generated through lentiviral transduction. Lentiviruses were produced by transfecting 293T cells with an equal amount of lentiviral packaging combo (pMDLg/pRRE (Addgene Plasmid 12251), pRSV-Rev (Addgene Plasmid 12253), VSV-G (pMD2.G, Addgene Plasmid 12259) in 1:1:1 molar ratio), and plasmid of interest, followed by a change of media of the to-be-infected cell lines at 12 hours post-transfection. The supernatant of 293T cells containing fresh lentivirus was collected every 24 hours after media change and applied to cells to be infected. Infected cells are selected with the appropriate antibiotics or sorted by flow cytometry based on the introduced fluorescent protein markers.

Generation of the D274A mutation of POLG in ARPE-19 cells was performed using CRISPR/Cas9 RNP delivery as previously described ^26^ using a crRNA (spacer: 5’-CCTGATATGAGCTCGGTCAA-3’) and a single-stranded donor (ssODN, YF467) containing the desired mutation and mutated PAM site. 24 homozygous clones were individually confirmed by genotyping PCR and Sanger sequencing and mixed in equal ratios.

The *MGME1* knock-out ARPE-19 cell pool used in Figure 1D and S2D were generated by CRISPR RNP delivery via nucleofection as in ^26^ using a crRNA targeting the exon 1 of *MGME1* gene (spacer: 5’-CTTCCGGCCACATGAGTAAG-3’). Nucleofected cells were recovered and expanded for a week. Knock-out efficiency of this cell pool was confirmed by the level of MGME1 protein via western blot analysis using MGME1 antibody (Proteintech 23178-1-AP). The *MGME1^-/-^*ARPE-19 cells used in Figure 5C-D were generated by CRISPR RNA delivery via nucleofection but with differences in crRNA (spacer: 5’-AGTTATTTCAGACCATTTGC-3’) and nucleofection condition (P3 solution and EA-104 program). Cells were harvested 48 hours post nucleofection and plated in clonal density. Single clones were manually picked, plated to 96-well plates, and genotyped by PCR using primers (Forward: 5’-GCACACGCCTCCCTTCCTG-3’; Reverse: 5’-GTCTGGGCGACGCCTCTG-3’) to screen for insertions and deletions. Four homozygous clones confirmed by western blots using MGME1 antibody (Sigma HPA0040913) were chosen, mixed in 1:1 ratio, and expanded for downstream experiments.

#### Subcellular fractionation

Cells were harvested and processed using Cell Fractionation Kit (Abcam ab109719) following standard protocol with optimization for ARPE-19 cells. 50 μl 1x Buffer A and 50 μl Buffer B (or Buffer C) were used for 500,000 cells. Buffer B was prepared by diluting Detergent I 500-fold in 1x Buffer A. Buffer C was prepared by diluting Detergent II 12.5-fold in 1x Buffer A. Cytosol and mitochondria extractions were both performed on a rotator for 30 min at RT. All centrifugation steps during the extraction were performed for 5 min at 4°C. Cytosolic, mitochondrial, and nuclear fractions were mixed with 5X SDS-PAGE Sample Buffer and shaken at 60°C for 10 min before western blot analysis.

#### Western blot analysis

For western blots of mtDNA^ΔScaI^ clones and in particular when analyzing mtDNA-encoded OXPHOS proteins, cells harvested were lysed in RIPA buffer (150 mM sodium chloride, 1% NP-40, 0.5% sodium deoxycholate, 0.1% SDS, 50 mM Tris, pH 8.0) and sonicated at 4°C. Protein concentration was measured using the Pierce BCA protein assay kit (Thermo Fisher Scientific) and equal micrograms of proteins were mixed with Laemmli buffer before being loaded to SDS-PAGE gels. For other western blots, cells harvested were counted and lysed in 10 μl Laemmli buffer (4% (w/v) SDS, 20% glycerol, 120 mM Tris-HCl, pH6.8) per 100,000 cells. Lysates were vortexed and boiled at 95°C for 10 min. Proteins from equivalent numbers of cells were loaded on SDS-PAGE gels. SDS-PAGE gels were transferred to nitrocellulose or PVDF membranes using Trans-Blot Turbo Transfer System or Mini Trans-blot cells (Bio-Rad). Membranes were blocked in 5% milk in TBST (137 mM NaCl, 2.7 mM KCl, 19 mM Tris Base, 0.1% Tween-20), followed by incubation with primary antibodies in 1% milk (1% BSA for p-eIF2α) overnight at 4°C. Subsequently, membranes were washed with TBST 3 times, incubated with HRP-linked secondary antibodies at 1:2,500 dilution, and washed again. Membranes were then developed with Clarity Western ECL Substrate or Clarity Max Western ECL Substrate (Bio-Rad) and imaged on ChemiDoc XRS+ Imager (Bio-Rad). Loading control a Rhodamine detected GAPDH conjugated antibody against GAPDH. Western blots were quantified using Image lab software (Bio-Rad).

#### Immunofluorescence

ARPE-19 cells stably expressing mito-Ku, mito-LigD, or mito-T4 alone were plated on coverslips in 6-well plates. Cells were stained with 125 nM MitoTracker Deep Red (Invitrogen) for 30 min. Cells were rinsed with PBS buffer and fixed with 4% paraformaldehyde for 10 min at RT. After PBS washes, cells were permeabilized with 0.5% Triton X-100 in PBS buffer for 10 min at RT, followed by PBS washes. Cells on coverslips were blocked with blocking buffer (1 mg/ml BSA, 3% goat serum, 0.1% Triton X-100, 1 mM EDTA) for 30 min at RT and incubated with Myc antibody (Cell Signaling Technology 2276) for mito-Ku, or FLAG antibody (Sigma F3165) for mito-LigD and mito-T4 in blocking buffer overnight at 4°C. Cells were then washed with PBS and hybridized with Alexa Fluor 488 conjugated anti-mouse secondary antibody for 1 hour at RT. After PBS washes and DAPI staining, coverslips were mounted on glass slides with ProLong Gold Antifade mountant (Invitrogen). Slides were imaged on a Nikon ECLIPSE Ei2-E inverted microscope (for mito-Ku and mito-LigD) or Nikon CSU-W1 Spinning Disk Confocal microscope (for mito-T4).

#### Genomic DNA purification

Total genomic DNA was purified for mtDNA deletion detection, measurements of heteroplasmy and copy number by qPCR, and southern blot analysis. Cell pellets harvested were resuspended in 400 μl PBS buffer containing 0.2% (w/v) SDS, 5 mM EDTA, and 0.2 mg/ml Proteinase K and incubated at 50°C for 6 hours with constant shaking at 1,000 rpm. DNA was precipitated by adding 0.3 M Sodium Acetate, pH 5.2, and 600 μl isopropanol and kept at −20°C for over 2 hours. Precipitated DNA was centrifuged at 20,000 x g at 4°C for 30 min, followed by a 70% ethanol wash. DNA was then resuspended in TE buffer (10 mM Tris-HCl, pH8.0, 0.1 mM EDTA) and quantified by Nanodrop.

#### Southern blot analysis

Purified genomic DNA was digested with BamHI-HF restriction enzyme (NEB) at 5 units per 1 μg DNA for 6 hours at 37°C. Products and Quick-Load 1kb Extend DNA Ladder (NEB) were separated on a 0.6% agarose gel supplemented with ethidium bromide at 30 V for 24 hours at 4°C. The ladders were imaged on ChemiDoc XRS+ Imager (Bio-Rad) before blotting. The agarose gels went through the following washes: 1) depurination (0.25 M HCl) for 30 min, 2) twice denaturation (1.5 M NaCl, 0.5 M NaOH) for 30 min, 3) twice neutralization (3M NaCl, 0.5 M Tris-HCl, pH7.0) for 30 min. The gels were blotted to Amersham Hybond-N membranes (GE) via overnight capillary transfer with 20x SSC (3 M NaCl, 0.3 M sodium citrate). The membranes were crosslinked under UV at 1200×100 μJ/cm^2^ and then incubated in pre-hybridization buffer (1.5x SSC, 1% SDS, 7.5% dextran sulfate, and 125 μg/ml salmon sperm DNA) at 65°C for 1 hour. Probes specific to human mtDNA were PCR amplified from 3 regions as in ^66^, and probes specific for nuclear 18S ribosomal DNA were amplified using primers as in ^27^. For probe labeling, 75 ng of each of the purified PCR products was mixed with 1 μl Random Hexamers (50 μM, Invitrogen) in a total volume of 30 μl, which was then boiled at 95°C for 5 min and cooled to room temperature. The reaction was supplemented with 1 μl DNA Polymerase I, Large (Klenow) fragment (NEB), 5 μl dCTP [α-32P] (3000Ci/mmol), 1x NEBuffer 2 and 33 μM dATP, dGTP, dTTP, and incubated at RT for 90 min. After adding 50 μl TNES buffer (20 mM EDTA, 400 mM NaCl, 0.5% SDS, 50 mM Tris base), reactions were deactivated at 65°C for 10 min. Probes were then purified through a Sephadex G-50 fine (GE) column made with TNE buffer (50 mM Tris-HCl, pH7.4, 100 mM NaCl, 0.1 mM EDTA) in a 3 ml syringe. Eluted probes (in 1 ml TNES buffer) were boiled at 95°C for 5 min and immediately added to the membrane in the pre-hybridization buffer. After overnight hybridization at 65°C, membranes were washed once with 4x SSC, three times with 2x SSC supplemented with 0.1% SDS, and wrapped and exposed to storage phosphor screens, which were scanned using Typhoon Imager (GE).

#### mtDNA deletion quantified by Quantitative PCR (qPCR)

Regions of interest in mtDNA and the nuclear ACTB locus were independently amplified using specific primers from total genomic DNA with SsoAdvanced SYBR Green Supermix (Bio-Rad). mtDNA^ΔScaI^ was amplified by YF174 and YF391 primers across deletion junctions. A non-deleted region (MT-RNR1/2) was amplified by YF338 and YF339 primers to indicate total mtDNA. A region across two ScaI sites spanning both deleted and non-deleted regions was amplified by YF391 and YF780 primers to indicate wild-type mtDNA. ACTB was amplified by YF454 and YF455 primers. qPCR reactions were in a volume of 10 μl per well per technical replicate, with 250 nM primers and 12.5 ng template genomic DNA, and run with standard cycling conditions suggested by Bio-Rad on Roche LightCycler 480 or Applied Biosystems QuantStudio 6 Real-Time PCR System. Quantification was based on the threshold cycle number (Ct value) derived from real-time PCR amplification. mtDNA^ΔScaI^ level was calculated by normalizing mtDNA^ΔScaI^ to ACTB. Total mtDNA level was calculated by normalizing mtDNA^Total^ to ACTB. mtDNA^ΔScaI^ heteroplasmy = (2^-(Ct^ΔScaI^) / (2^-(Ct^ΔScaI^) + (2^-(Ct^WT^)) * 100%.

#### mtDNA deletion detection by junction PCR and agarose gel electrophoresis

To detect formation of mtDNA^ΔScaI^ after inducing mito-ScaI cleavage in cells expressing mito-EJ components, genomic DNA was purified and amplified using Q5 High-Fidelity DNA Polymerase (NEB M0491L) and primers (YF176 and YF411) ∼1 kb upstream and downstream of the most distal ScaI sites. PCR products were run on a 1.2% agarose gel in 0.5 x Tris-Borate-EDTA (TBE) Buffer. Direct ligation between the two most distal ScaI sites will yield a 2003 bp PCR product. Due to the mitochondrial exonuclease activity on linear mtDNA, the repaired mtDNA formed in a pool of mito-EJ cells upon ScaI cleavage carried extended deleted regions from the ligation junction, yielding smaller PCR products that were presented as a smear on agarose gel.

#### Assay for Transposase-Accessible Chromatin using sequencing (ATAC-seq) and mtDNA deletion identification

For Figure 1C and S2A, ARPE-19 cells that were wild-type, stably co-expressing mito-Ku and mito-LigD, or expressing mito-T4 DNA Ligase alone were infected with doxycycline (Dox)-inducible mito-ScaI (iScaI, TLCV2-ScaI-T2A-EGFP-Puro) lentivirus to generate three iScaI cell lines with and without mito-EJ. iScaI cell lines were treated with 1 μg/ml doxycycline for 12 hours to induce ScaI cleavage in mtDNA, followed by doxycycline removal and recovery for five days. Recovered cells and cells without doxycycline treatment were harvested for ATAC-seq. For Figure 1D and S2D, wild-type, *MGME1* knock-out, and *POLG^D274A/D274A^* ARPE-19 cells were infected with mito-T4 DNA ligase lentivirus (produced from plasmid pHAGE-EF1a-mito-T4-IRES-tdTomato) to generate three mito-T4 expressing cell lines with different genetic backgrounds. These three cell lines were transfected with mito-ScaI plasmid (pcDNA-mitoScaI) by nucleofection to induce ScaI cleavage in mtDNA. Cells were harvested three days post-transfection for ATAC-seq.

For Figures 5C and D, *MGME1^-/-^* ARPE-19 cells were infected with Dox-inducible mito-ApaLI (iApaLI) lentivirus (produced from plasmid TLCV2-ApaLI-T2A-EGFP-Puro) to generate iApaLI cell line. *MGME^-^*^/-^ iApaLI cell lines were subsequently infected with mito-Ku and mito-LigD lentiviruses (produced from plasmid pHAGE-EF1a-mito-Ku-IRES-Hygro and pHAGE-EF1a-mito-LigD-IRES-tdTomato, respectively), followed by selection with 300 μg/ml Hygromycin B (Thermo Fisher Scientific 10687010) and fluorescence-activated cell sorting (FACS) to enrich for Ku and LigD co-expressing cells. *MGME^-/-^*iApaLI cell lines with and without mito-EJ components were treated with 1 μg/ml doxycycline for 12 hours to induce ApaLI cleavage in mtDNA, followed by doxycycline removal and recovery for 5 days. Recovered cells and cells without doxycycline treatment were harvested for ATAC-seq.

50,000 cells were collected and frozen in 10% DMSO in culture media in −80°C before submission to the Integrated Genomics Operation (IGO) core at Memorial Sloan Kettering Cancer Center. Library preparation followed standard ATAC-seq protocol without nuclei isolation^67^. Deletions and duplications were called using MitoSAlt^34^, a pipeline that classifies deletions that overlap with replication origins as duplications with the assumption that mtDNA deletions with altered replication origins are unlikely to be propogated. We applied the default filtering criteria: cluster threshold = 5, break threshold = −2, hplimit = 0.01, deletion threshold min = 30, breakspan = 15, split distance threshold = 5, score threshold = 80, and evalue threshold = 0.00001.

One exception is for Figure 5C-D, where ApaLI was used to generate a single DNA break at the MT-RNR1 (∼1.6 kb away from the replication origin OriH). We expected that the scale of deletions would be smaller and may intersect with the replication origin OriH. Therefore, we made the following changes to accommodate the analysis: 1) deletion threshold min = 10; 2) disable the reclassification of deletions into duplications, i.e. classify all events as deletions by setting no replication origins (orihs = −1, orihe = −1, orils = −1, orile = −1). Moreover, we predicted that suppression of linear DNA degradation due to loss of MGME1 may result in small insertions and deletions (indels) at the ApaLI site in the presence of mito-EJ. Therefore, we called mitochondrial DNA variants in the ATAC-seq data as previously described in ^68^. Briefly, variant calls from Mutect2 intersected with those from our in-house variant caller, which utilizes the SAMtools mpileup. Variants located in the black-listed regions (513–525 and 3105–3109) were subsequently filtered out. Variants across all the samples were aggregated and genotyped in individual samples to calculate heteroplasmy levels. All variants within the ApaLI site (1586-1591) were analyzed and plotted by their heteroplasmy levels.

#### Isolation of mtDNA^ΔScaI^ clonal cells

A founder ARPE-19 cell line stably expressing mito-T4 DNA ligase (pHAGE-EF1a-mito-T4-IRES-tdTomato) and Dox-inducible mito-ScaI (TLCV2-ScaI-T2A-EGFP-Puro) was used to isolate clonal cells carrying varying levels of mtDNA^ΔScaI^. To obtain individual clonal cells, the founder cells were sorted into 96-well plates with one cell per well. Clonally expanded cells were split into two plates with one plate for mtDNA^ΔScaI^ genotyping and the other for maintenance. For the genotyping plate, cells in each well were lysed in 50 µl lysis buffer (1x PBS, 10mM Tris HCl pH 8, 0.5% NP-40, 0.5% Tween20, freshly supplemented with 200 µg/ml Proteinase K) at 50°C overnight followed by inactivation at 95°C for 10 min. 1 µl of crude lysate was amplified in a 10 µl reaction containing 5 µl 2x FailSafe PCR Buffer H (Lucigen FSP995H), 0.1 µl Taq Polymerase (NEB M0273), and 250 nM genotyping primers (YF391 and YF174). While wild-type mtDNA was either a 3938 bp band or not present due to short PCR extension time, one or more bands below 491 bp indicated the presence of deletions. About 10% of clones derived from the founder cells carried detectable mtDNA^ΔScaI^. Notably, activating robust mito-ScaI expression by Dox treatment was unnecessary to obtain clones carrying intermediate mtDNA^ΔScaI^. On the contrary, induction of robust ScaI cleavage by Dox led to mtDNA^ΔScaI^ cells that were all homoplasmic. This suggests that deletions may have been formed upon transduction of T4 and ScaI, or through the basal expression of ScaI. To reduce the residual expression of ScaI, we used CRISPR (gRNA spacers: 5’-TCAACGATCAGCTTCCCCGG-3’ and 5’-GAGTTGAGTGTTGGATACGA-3’) to target the transgene ScaI in the founder cell line and sorted out GFP negative cells after Dox induction, considering that a frameshift in ScaI would lead to a frameshift in subsequent T2A-EGFP. 8 out of 10 selected clones for in-depth characterization were derived after ScaI targeting.

Clones selected from genotyping PCR were further examined for the number of deletion types, deletion junctions, and heteroplasmy levels. We used primers further away from the distal ScaI sites to probe for larger deletions (Figure S3B) and chose clones with a single deletion. PCR products were sequenced by Sanger sequencing to determine the ligation junction. Despite the variable extent of degradation leading to different deletion junctions in clones, they were all deleted from within COX2 to within ND4 (Figure S3D). Several clones with no detectable mtDNA^ΔScaI^ in genotyping PCR were randomly chosen to be the 0% clones.

#### mtDNA fluorescent in-situ hybridization (FISH) and quantification

FISH probes were designed based on bulk ATAC-seq analysis results. Regions consistently deleted or never deleted across all identified deletions were selected as probe-binding regions. To generate probe DNA, 100 ng of purified genomic DNA of ARPE-19 cells was amplified using Advantage 2 Polymerase Mix (TaKaRa 639201) in 50 μl reactions. The deleted probe was amplified using primers YF748 and YF749 with 57°C as the annealing temperature and 3 min extension for 35 cycles, and the non-deleted probe was amplified using primers YF743 and YF744 with 56°C as the annealing temperature and 3 min extension for 35 cycles. PCR products were resolved on an agarose gel and gel purified to obtain at least 80 ng/μl concentration of DNA. The two DNA probes were subsequently labeled with different fluorescent dyes via nick translation using FISH Tag™ DNA Multicolor Kit, Alexa Fluor™ dye combination (Thermo Fisher Scientific F32951) following manufacturer’s instructions.

mtDNA FISH was performed as described previously ^69^ with minor modifications. Cells grew on coverslips (Fisher scientific 50-949-008) were rinsed by PBS and fixed in fixation buffer (5% paraformaldehyde (PFA, Electron Microscopy Sciences 15713), 100 mM sodium cacodylate pH 7.2, 100 mM sucrose, 40 mM potassium acetate, 20 mM sodium acetate, 10mM EGTA) for 4 min at room temperature (RT). Cells on coverslips were washed once in 2x SSCT (2x SSC, 0.1% Tween) for 10 min and incubated in 100 μg/ml RNase in 2x SSCT for 30 min, followed by a wash in 2x SSCT for 10 min at RT. Cells were further incubated for 10 min in 40% formamide (Thermo Fisher Scientific 4311320) in 2x SSCT, 10 min in 20% formamide in 2x SSCT, and 10 min in 50% formamide in 2x SSCT twice at RT. Coverslips were subsequently incubated in hybridization mix (50% formamide, 2x SSC, 10% dextran sulfate, 0.125 μg/ul E coli. tRNA, 0.125 μg/ul salmon sperm DNA, 50 ng (0.5 μl) of each fluorescent probe) on a glass slide, sealed with Ctyobond Removable Coverslip Sealant (SciGene 2020-00-1), denatured at 91°C for 3 min, and incubated at 37°C overnight. After removing the sealant, coverslips went through a series of washes: 10 min in 50% formamide in 2x SSCT at 37°C for three times, 10 min in 40% formamide in 2x SSCT at RT, 20% formamide in 2x SSCT at RT, 10 min in 2x SSCT at RT for three times, and 10 min in PBS at RT for three times. Coverslips could either be re-fixed with 2% PFA in PBS for 30 min for standard immunofluorescence protocol as mentioned above, or stained with DAPI in PBS and mounted on glass slides using Prolong Glass Antifade Mountant (Thermo Fisher Scientific P36980).

Cells were imaged on the Nikon Eclipse Ti2 inverted microscope with 60x oil DIC lens and the Yokogawa CSU-W1 SoRa confocal scanner unit at SoRa mode and 4x magnification. Z-stacks were captured at 0.15 μm step size for ∼17 steps. The foci count of deleted and non-deletion regions per cell were quantified using a custom ImageJ script on Fiji software. In brief, individual cells were outlined manually as regions of interest (ROIs), followed by segmentation using the Labkit – intuitive pixel classification Fiji plugin. Segmented images were then processed by Z projection by max intensity, “Convert to mask,” and “Watershed.” While the foci of the deleted probe indicate the wild-type mtDNA, the foci of the non-deleted probe indicate the total mtDNA. Thus, the segmentation of deleted mtDNA was obtained using the “Image Calculator” function to subtract the segmentation of wild-type mtDNA from that of total mtDNA. Wild-type mtDNA foci were counted by the “Analyze Particles” function (size = 6-Inifinity, circularity = 0.3-1), and deleted mtDNA foci were counted using a more stringent circularity (0.85-1). Heteroplasmy = deletion counts / (deletion counts + foci counts) * 100%.

#### Mitochondrial network analysis

mtDNA^ΔScaI^ clonal cells cultured in Tet-free serum were stained with 250 nM abberior LIVE ORANGE and 5 μg/mL Hoechst 33342 (NucBlue Live ReadyProbes Reagent) for 45 minutes. Stained cells were rinsed twice with PBS and cultured in phenol-red free for live-cell imaging. Cells were imaged on the Nikon Eclipse Ti2 inverted microscope with 60x oil 1.49 NA DIC lens and the Yokogawa CSU-W1 SoRa confocal scanner unit at SoRa mode and 2.8x magnification. Z-stacks were captured at 1.63 μm step size for ∼30 steps. Images were deconvoluted using NIS-Elements software.

#### Mitochondrial membrane potential and apoptosis analysis

mtDNA^ΔScaI^ clonal cells cultured in Tet-free medium were stained with 1 μM Tetramethylrhodamine, Methyl Ester, Perchlorate (TMRM, Thermo Fisher Scientific T668) in culture media for 30 min at 37°C. Cells were then harvested by trypsinization, centrifuged, resuspended in 3% FBS in PBS, and analyzed on an LSR Fortessa analyzer. TMRM signal was detected through a 561 nm laser and 586/15 Bandpass filter. Recorded data were analyzed using FlowJo software. As a positive control, 0% clones were treated with 20 μM CCCP for 6 hours.

For apoptosis analysis by Annexin V staining, mtDNA^ΔScaI^ clonal cells were seeded in equal cell numbers in a Tet-free medium. The next day, cells were rinsed with PBS and changed to glucose or galactose medium. Because the near homoplasmic mtDNA^ΔScaI^ cells underwent cell death within 24 hours in galactose medium, we harvested cells 24 hours after media change. A culture medium containing floating dead cells was collected together with trypsinized cells. Cells were stained using Pacific Blue Annexin V Apoptosis Detection Kit with 7-AAD (BioLegend 640926) per the manufacturer’s instructions. Cells were analyzed on an LSR Fortessa analyzer. Pacific Blue Annexin V signal was detected through a 405 nm laser and 450/50 Bandpass filter. Recorded data were analyzed using FlowJo software.

#### Oxygen consumption measurement

Oxygen consumption rate (OCR) was measured using an XFe96 Extracellular Flux Analyzer (Agilent) following the manufacturer’s instructions. Cells were plated in Seahorse microplates (Agilent) at 20,000 cells per well. After overnight, when cells adhered to the plates, cell culture media were removed and replaced with Seahorse media (DMEM, Agilent 103575) supplemented with 10 mM glucose and 4 mM glutamine. OCR analysis was performed at basal conditions and after injections of oligomycin (1 μM), FCCP (1 μM), and rotenone plus antimycin A mix (each at 1 μM), where indicated according to the manufacturer’s instructions.

#### Metabolite analysis using GC-MS

All cells were seeded in 6-well plates 48 hours prior to harvest and washed once with PBS prior to the medium change. Regular DMEM containing 25mM glucose and 4mM glutamine supplemented with 10% dialyzed FBS, 1% Penicillin-Streptomycin and 50 μg/ml μM uridine. 1 mM pyruvate was used when indicated. For comparing metabolite levels in cells cultured in media with or without pyruvate, the media were replaced 24 hours after seeding, and the cells were cultured in the indicated media for 24 hours. Metabolism was quenched with 1 ml ice-cold 80% methanol containing 2 μM deuterated 2-hydroxyglutarate (d-2-hydroxyglutaric-2,3,3,4,4-d5 acid (d5-2HG)) and stored at −80°C overnight. The methanol-extracted metabolites were collected and centrifuged at 21,000 x g for 30 min to remove proteins. The supernatant was dried in a vacuum evaporator (Genevac EZ-2 Elite) for 6 hours. Dried metabolites were dissolved in 20 mg/mL methoxyamine hydrochloride (Sigma 226904) in pyridine (Thermo Fisher Scientific TS-27530) for 90 min at 30°C and derivatized with MSTFA containing 1% TMCS (Thermo Fisher Scientific TS-48915) for 30 min at 37°C. Samples were analyzed using an Agilent 7890A GC connected to an Agilent 5975C Mass Selective Detector with electron impact ionization. The GC was operated in splitless mode with constant helium gas flow at 1 mL/min. 1 μL of derivatized metabolites was injected onto an HP-5MS column (15 m x 0.25 mm, 0.25 μm particle size), the inlet temperature was 250°C, and the GC 6 oven temperature was ramped from 60 to 290°C over 25 min. Data analysis was performed using Mass Hunter Quantitative Analysis software v10.0 (Agilent). Peaks were normalized to the surrogate internal standard (d5-2HG) peak area and biomass (cell number x average cell size). Ions used for quantification of steady-state metabolite levels are as follows: d5-2HG m/z 354; citrate m/z 273; malate m/z 233; glutamate m/z 363; aspartate m/z 232. All peaks were manually inspected and verified relative to known spectra for each metabolite.

#### Growth analysis by Incucyte

Clonal cells were seeded in Tet-free medium in 48-well plates with 7,000 cells per well and three wells per experimental condition. Over 12 hours later, when cells were fully attached, cells were rinsed with PBS and replaced with treatment media (-Pyruvate medium was DMEM containing 25 mM glucose and 4 mM glutamine and supplemented with 10% dialyzed FBS, 1% Penicillin-Streptomycin, and 50 μg/ml μM uridine; + Pyruvate medium was - Pyruvate medium supplemented with 1 mM sodium pyruvate). Plates were imaged every 12 hours on the Incucyte S3 Live-cell analysis system (Sartorius) using brightfield and 10x objective. 16 fields per well were captured. Cell confluency was measured using Incucyte analysis software with settings as follows: 0.5 Segmentation Adjustment, 700 μm2 Hole Fill, 2000 μm2 minimal area.

#### Modified DOGMAseq (High LLL DOGMAseq)

Single Cell Multiome ATAC + Gene Expression was performed with 10X genomics system using Chromium Next GEM Single Cell Multiome Reagent Kit A (catalog no. 1000282) and ATAC Kit A (catalog no. 1000280). We followed Chromium Next GEM Single Cell Multiome ATAC + Gene Expression Reagent Kits User Guide with the difference that single cell suspension was prepared almost as described in Mimitou et al., ^28^ (DOGMAseq LLL protocol with NP-40 buffer). However, to further increase mitochondria DNA coverage, we optimized the lysis step by increasing NP-40 concentration to 0.3% and the time of lysis to 3’. Final libraries were sequenced on Illumina NovaSeq X+ platform (R1 – 28 cycles, i7 – 8 cycles, R2 – 90 cycles).

In addition, in clones with 0, 25, 40, 41, 58, 60, 75, and 100% heteroplasmy, samples were multiplexed together on one lane of 10X Chromium (using Hash Tag Oligonucleotides - HTO) following previously published protocol^70^ (Figure S5A,B). Final libraries were sequenced on Illumina NovaSeq X+ platform (R1–28 cycles,i7–8 cycles,R2–90cycles).

#### Computational methods

##### DOGMAseq preprocessing

All DOGMAseq data presented in this manuscript were processed using Cell Ranger ARC, which uses Illumina fastq files as input and performs alignment and PCR duplicates and repeat ATAC reads removal to create an output for a gene expression count matrix and ATAC fragments file. CellRanger ARC reference was built after the blacklist, and the chrM contig was moved from “non-nuclear” to “nuclear.” This ensured the mito fragments were not discarded by CellRanger and appeared in the fragments file. We utilized a modified hg38 reference genome to enrich for reads from mtDNA for which we blacklisted regions of the nuclear genome that share high homology with mtDNA using the hg38 bedfile from (https://github.com/caleblareau/mitoblacklist) to recover otherwise multi-mapped mtDNA reads^71^. The ATAC fragment file records each sequenced fragment’s cell barcode, chromosomal location, and start and end locations.

##### Processing scRNA-seq

We used the default filtered matrix from Cell Ranger ARC to filter for high-quality cells. All downstream processing steps were taken using Scanpy preprocessing functions. First, we processed scRNA-seq data of all three conditions (Galactose, Glucose + Pyruvate, and Glucose - Pyruvate) together. We performed median library size normalization and log normalization and ran PCA (npca = 30) using 4500 highly variable genes. Next, we clustered the cells using the Leiden algorithm (resolution parameter = 0.1). We identified a unique cluster of low-quality cells with greater than 20% counts from ribosomal genes and less than 4500 total gene counts. We subsequently removed this population of low-quality cells, resulting in 25,509 high-quality cells (6787 Galactose, 7817 Glucose + Pyruvate, 10905 Glucose - Pyruvate). In the case of the eight-clone mix (Figure S5), we used the default filtered matrix from Cell Ranger ARC to filter for high-quality cells. We also filtered for cells with an average mitochondrial genome coverage of at least 20 (see *Mitochondrial genome coverage* section). We used the resulting normalized count matrix of high-quality cells for additional down-stream analysis described below.

After filtering out low-quality cells, we reprocessed the Galactose, Glucose + Pyruvate, and Glucose – Pyruvate samples altogether starting from the raw counts. Again, we first performed median library size normalization, log normalization. Next, we performed PCA on the 13 mitochondrial genes. Then, we computed nearest neighbors using sc.pp.neighbors(n = 30), and plotted using sc.tl.umap(min_dist = 0.1) (Figure 3D, E). We also computed nearest neighbors directly on the log-normalized counts of mitochondrial genes instead of PCs, and the results were robust.

##### Mitochondrial genome coverage

To compute single-cell mitochondrial genome coverage, we filtered the ATAC fragments CSV output from Cell Ranger ARC for fragments from mtDNA and grouped them by cell barcodes. Next, for each cell, we aggregated all mapped ATAC fragments and obtained the coverage per base pair along the mitochondrial genome. Specifically, we define coverage per base pair (or coverage per nucleotide) as the # of distinct reads that overlap the given base pair within a single cell. Next, we compute an overall coverage across the mitochondrial genome by averaging the coverage per base pair from chrM:0 – 16569. For all experiments, we filtered for cells with at least 20 average coverage per nucleotide across the mitochondrial genome.

We observed many mitochondrial fragments with lengths greater than 3000 bp that spanned the deleted region. We reasoned this is due to the paired-end nature of sequencing, where only up to 50 bp is captured from either end of the fragments. Indeed, ATAC-seq reads are typically expected to be ∼150 bp ^67,72^ and certainly less than 1000 bp in a standard sequencing platform, suggesting such large fragments represent mitochondrial genomes with successful deletion. As such, to accommodate these fragments, we split each into two with 50 bp padding from the start and end. For example, a mitochondrial read spanning base pairs 7000-11000 is split into the following reads: 7000 – 7050 and 10950 – 11000.

##### Estimating Heteroplasmy

mtDNA heteroplasmy at a single-cell level was estimated based on the coverage of a deleted region and a common region in the scATAC-seq data. The deleted region (8013-11460) is between the two most distal ScaI sites. The common region (1231-4678), in the same length as the deleted region, was selected because it was not deleted based on the bulk ATAC-seq results.

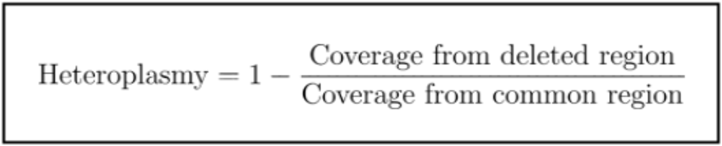

Heteroplasmy is then computed as one minus the ratio of the number of fragments that map to the deleted region to the number of fragments that map to the undeleted common region. Higher heteroplasmy results from low coverage in the deleted region, which signifies more deletions. In cases with more coverage from the deleted region than the common region, we classify these cells as 0% heteroplasmy.

##### Identification of genes correlating with heteroplasmy

We first computed differential gene expression analysis between discrete heteroplasmy levels in Galactose, Glucose + Pyruvate, and Glucose – Pyruvate samples separately. We defined the following categories based on heteroplasmy: Zero [0 – 0.05), Low [0.05 – 0.4), Medium [0.4 – 0.75), High [0.75, 1]. For each sample, we performed median library size normalization and log normalization. Next, we report the differentially expressed genes between pairwise Low vs Zero, Medium vs Zero, and High vs Zero heteroplasmy cells respectively using Scanpy’s *sc.tl.rank_gene_groups(reference = ‘Zero’, method = ‘Wilcoxon’)* (Fig 4A).

Next, for each sample (Galactose, Glucose + Pyruvate, and Glucose - Pyruvate), we searched for genes that correlate with heteroplasmy in a continuous manner (Fig 3F, 4D-E S7-D). Note this analysis is done independent of the discrete analysis. The sparsity and noise of scRNA-seq data can impact computing correlation at the single-cell level. To circumvent this issue, we applied a few noise mitigation techniques. First, we removed genes that were expressed in less than ten cells Then, we computed correlation using two independent approaches (described below) and obtained highly concordant results using both approaches.

###### Method 1: Averaging cells based on heteroplasmy

To account for noise of scRNA-seq data, we sorted cells by heteroplasmy and averaged the gene expression raw counts for every twenty consecutive cells. Thus, for a group of twenty cells with similar heteroplasmy, we obtain a more robust “average” cell. Note the heteroplasmy of the “average” cell is the average heteroplasmy of the cells in the group. Using these “averaged” cells, we then performed library size normalization using Scanpy’s *sc.pp.normalize_total()* followed by log normalization. This method was preferred over just averaging cells within a heteorplasmy bin (e.g. [0 – 0.05, 0.05 – 1, …, 0.95 – 1]) because it is unaffected by the uneven distribution of cells across heteroplasmy (Figure 3B).

###### Method 2: MAGIC-based imputation

Independently, we applied MAGIC^73^, a graph-diffusion based imputation algorithm that removes sparsity and noise in single-cell gene expression data. It has been successfully used in various biological contexts in the field^74–76^. Since we are interested in dynamics driven by deletion in the mtDNA, we chose to apply MAGIC imputation (Knn = 30, t = 3) on the nearest neighbor graph constructed using only mitochondrial genes as features (post library size and log normalization). The choice of only mitochondrial genes as features is motivated by our goal of correlating cells by heteroplasmy and our observation that variation in our single-cell data is almost entirely driven by mitochondrial gene expression.

Based on both heteroplasmy-binned and MAGIC-imputed expression, we computed the Pearson and Spearman correlations, as well as the slopes of defined heteroplasmy windows for each gene’s expression relative to heteroplasmy.

###### Filtering criteria to distinguish “heteroplasmy-sensing” vs. “threshold-triggered” genes

We initially filtered for genes with at least 10 total counts using *sc.pp.filter_genes(min_counts=10)*. Next, to identify gene expression dynamics as a function of heteroplasmy, including both threshold-triggered and heteroplasmy-sensing genes, we applied a three-step gene filtering. First, for each gene, we computed the Spearman correlation between heteroplasmy and MAGIC imputed expression and identified genes that had Spearman correlation > 0.3 and < −0.3. This cutoff roughly includes the top 10% and bottom 10% of genes. Second, for the genes identified in the first step, we binned the heteroplasmy into two groups with 0.75 as the cut-off based on the metabolic threshold we identified in Figure 2. We determined the slope of each group (heteroplasmy ranges 0 – 0.75 and 0.75 – 1) and computed the log2 fold change of the slopes as follows: log2[abs(slope^0.75–1^)/ abs(slope^0-0.75^)]. Genes with a log2FC >= 2 were classified as threshold-triggered, and genes with a −2 < log2FC < 2 were classified as heteroplasmy-sensing. Finally, for our third filtering step, we filtered threshold-triggered genes for abs(slope^0.75–1^) > 0.1 and filtered heteroplasmy-sensing genes for abs(slope^0-0.75^) > 0.05 or abs(slope^0.75–1^) > 0.05. These thresholds roughly resulted in 25% of the threshold-triggered and 15% of heteroplasmy-sensing genes that passed the second filtering step. The final filter allowed us to select genes with a large absolute effect size since log2FC only captures relative change.

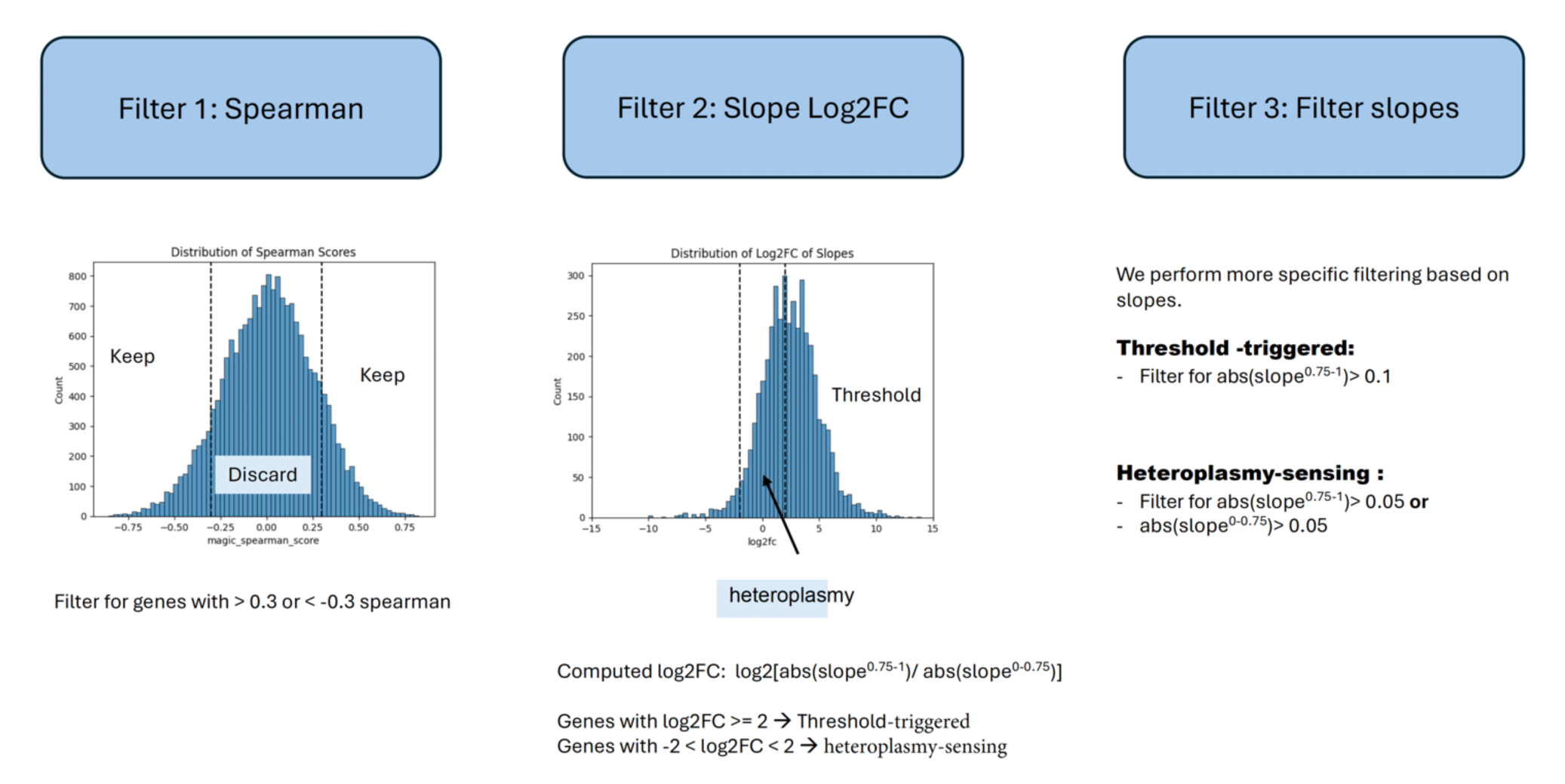

##### Estimation of mtDNA content from DOGMAseq

We adapted a previously proposed method originally developed to estimate mtDNA data from the bulk-level analysis ^77,78^ for computing mtDNA content from single-cell data. mtDNA content was estimated from ATAC-seq reads by applying a normalizing factor derived from coverage in the accessible regions of the nuclear DNA. A main challenge in this comparison arises from the difference in accessibility between nuclear DNA and mtDNA using scATAC-seq protocol with Tn5 transposase. Due to the heterochromatin nature of nuclear DNA, it is largely inaccessible, with ∼97% of the regions being closed. In contrast, mtDNA, which lacks nucleosomes, is uniformly accessible. To circumvent this issue, we focused on the most accessible regions of nuclear DNA. Specifically, we divided the genome 500bp tiles and calculated an accessibility score for each using the number of mapped fragments. We then selected the top 2.5% of tiles with the highest accessibility were selected. By assuming that mtDNA is sequenced at the same rate as these open regions in the nuclear DNA, we could accurately estimate mtDNA content using:

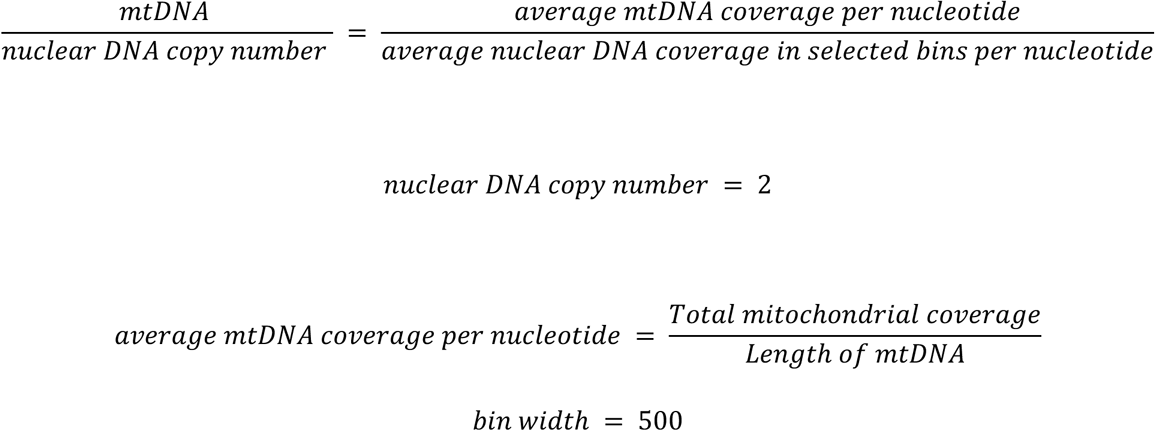

##### Pathway Analysis

To identify biological pathways enriched in upregulated and downregulated nuclear genes (Heteroplasmic_vs_WT_DEGs), in medium or high heteroplasmy in cell cultures in glucose + pyruvate, glucose – private, and galactose + pyruvate, we used the analysis tool at www.reactome.org. Data is provided in Table S3.

## REFERENCES

1. Holt, I.J., Harding, A.E., and Morgan-Hughes, J.A. (1988). Deletions of muscle mitochondrial DNA in patients with mitochondrial myopathies. Nature 331, 717–719. 10.1038/331717a0.

2. Moraes, C.T., DiMauro, S., Zeviani, M., Lombes, A., Shanske, S., Miranda, A.F., Nakase, H., Bonilla, E., Werneck, L.C., Servidei, S., and, et al. (1989). Mitochondrial DNA deletions in progressive external ophthalmoplegia and Kearns-Sayre syndrome. N Engl J Med 320, 1293–1299. 10.1056/NEJM198905183202001.

3. Schon, E.A., Rizzuto, R., Moraes, C.T., Nakase, H., Zeviani, M., and DiMauro, S. (1989). A direct repeat is a hotspot for large-scale deletion of human mitochondrial DNA. Science 244, 346–349. 10.1126/science.2711184.

4. Bua, E., Johnson, J., Herbst, A., Delong, B., McKenzie, D., Salamat, S., and Aiken, J.M. (2006). Mitochondrial DNA-deletion mutations accumulate intracellularly to detrimental levels in aged human skeletal muscle fibers. Am J Hum Genet 79, 469–480. 10.1086/507132.

5. Kraytsberg, Y., Kudryavtseva, E., McKee, A.C., Geula, C., Kowall, N.W., and Khrapko, K. (2006). Mitochondrial DNA deletions are abundant and cause functional impairment in aged human substantia nigra neurons. Nat Genet 38, 518–520. 10.1038/ng1778.

6. Tyynismaa, H., Mjosund, K.P., Wanrooij, S., Lappalainen, I., Ylikallio, E., Jalanko, A., Spelbrink, J.N., Paetau, A., and Suomalainen, A. (2005). Mutant mitochondrial helicase Twinkle causes multiple mtDNA deletions and a late-onset mitochondrial disease in mice. Proceedings of the National Academy of Sciences of the United States of America 102, 17687–17692. 10.1073/pnas.0505551102.

7. Spelbrink, J.N., Li, F.Y., Tiranti, V., Nikali, K., Yuan, Q.P., Tariq, M., Wanrooij, S., Garrido, N., Comi, G., Morandi, L., et al. (2001). Human mitochondrial DNA deletions associated with mutations in the gene encoding Twinkle, a phage T7 gene 4-like protein localized in mitochondria. Nat Genet 28, 223–231. 10.1038/90058.

8. Van Goethem, G., Dermaut, B., Lofgren, A., Martin, J.J., and Van Broeckhoven, C. (2001). Mutation of POLG is associated with progressive external ophthalmoplegia characterized by mtDNA deletions. Nat Genet 28, 211–212. 10.1038/90034.

9. Fu, Y., Tigano, M., and Sfeir, A. (2020). Safeguarding mitochondrial genomes in higher eukaryotes. Nature structural & molecular biology 27, 687–695. 10.1038/s41594-020-0474-9.

10. Gitschlag, B.L., Kirby, C.S., Samuels, D.C., Gangula, R.D., Mallal, S.A., and Patel, M.R. (2016). Homeostatic Responses Regulate Selfish Mitochondrial Genome Dynamics in C. elegans. Cell Metab 24, 91–103. 10.1016/j.cmet.2016.06.008.

11. Lin, Y.F., Schulz, A.M., Pellegrino, M.W., Lu, Y., Shaham, S., and Haynes, C.M. (2016). Maintenance and propagation of a deleterious mitochondrial genome by the mitochondrial unfolded protein response. Nature 533, 416–419. 10.1038/nature17989.

12. Yang, Q., Liu, P., Anderson, N.S., Shpilka, T., Du, Y., Naresh, N.U., Li, R., Zhu, L.J., Luk, K., Lavelle, J., et al. (2022). LONP-1 and ATFS-1 sustain deleterious heteroplasmy by promoting mtDNA replication in dysfunctional mitochondria. Nature cell biology 24, 181–193. 10.1038/s41556-021-00840-5.

13. Kandul, N.P., Zhang, T., Hay, B.A., and Guo, M. (2016). Selective removal of deletion-bearing mitochondrial DNA in heteroplasmic Drosophila. Nat Commun 7, 13100. 10.1038/ncomms13100.

14. Inoue, K., Nakada, K., Ogura, A., Isobe, K., Goto, Y., Nonaka, I., and Hayashi, J.I. (2000). Generation of mice with mitochondrial dysfunction by introducing mouse mtDNA carrying a deletion into zygotes. Nat Genet 26, 176–181. 10.1038/82826.

15. Tyynismaa, H., and Suomalainen, A. (2009). Mouse models of mitochondrial DNA defects and their relevance for human disease. EMBO Rep 10, 137–143. 10.1038/embor.2008.242.

16. Grady, J.P., Murphy, J.L., Blakely, E.L., Haller, R.G., Taylor, R.W., Turnbull, D.M., and Tuppen, H.A. (2014). Accurate measurement of mitochondrial DNA deletion level and copy number differences in human skeletal muscle. PloS one 9, e114462. 10.1371/journal.pone.0114462.

17. Lareau, C.A., Dubois, S.M., Buquicchio, F.A., Hsieh, Y.H., Garg, K., Kautz, P., Nitsch, L., Praktiknjo, S.D., Maschmeyer, P., Verboon, J.M., et al. (2023). Single-cell multi-omics of mitochondrial DNA disorders reveals dynamics of purifying selection across human immune cells. Nat Genet 55, 1198–1209. 10.1038/s41588-023-01433-8.

18. Hayashi, J., Ohta, S., Kikuchi, A., Takemitsu, M., Goto, Y., and Nonaka, I. (1991). Introduction of disease-related mitochondrial DNA deletions into HeLa cells lacking mitochondrial DNA results in mitochondrial dysfunction. Proceedings of the National Academy of Sciences of the United States of America 88, 10614–10618. 10.1073/pnas.88.23.10614.

19. Prigione, A., Lichtner, B., Kuhl, H., Struys, E.A., Wamelink, M., Lehrach, H., Ralser, M., Timmermann, B., and Adjaye, J. (2011). Human induced pluripotent stem cells harbor homoplasmic and heteroplasmic mitochondrial DNA mutations while maintaining human embryonic stem cell-like metabolic reprogramming. Stem Cells 29, 1338–1348. 10.1002/stem.683.

20. Cherry, A.B., Gagne, K.E., McLoughlin, E.M., Baccei, A., Gorman, B., Hartung, O., Miller, J.D., Zhang, J., Zon, R.L., Ince, T.A., et al. (2013). Induced pluripotent stem cells with a mitochondrial DNA deletion. Stem Cells 31, 1287–1297. 10.1002/stem.1354.

21. Gammage, P.A., Rorbach, J., Vincent, A.I., Rebar, E.J., and Minczuk, M. (2014). Mitochondrially targeted ZFNs for selective degradation of pathogenic mitochondrial genomes bearing large-scale deletions or point mutations. EMBO Mol Med 6, 458–466. 10.1002/emmm.201303672.

22. Bacman, S.R., Williams, S.L., Pinto, M., Peralta, S., and Moraes, C.T. (2013). Specific elimination of mutant mitochondrial genomes in patient-derived cells by mitoTALENs. Nat Med 19, 1111–1113. 10.1038/nm.3261.

23. Srivastava, S., and Moraes, C.T. (2001). Manipulating mitochondrial DNA heteroplasmy by a mitochondrially targeted restriction endonuclease. Hum Mol Genet 10, 3093–3099. 10.1093/hmg/10.26.3093.

24. Mok, B.Y., de Moraes, M.H., Zeng, J., Bosch, D.E., Kotrys, A.V., Raguram, A., Hsu, F., Radey, M.C., Peterson, S.B., Mootha, V.K., et al. (2020). A bacterial cytidine deaminase toxin enables CRISPR-free mitochondrial base editing. Nature 583, 631–637. 10.1038/s41586-020-2477-4.

25. Nissanka, N., Bacman, S.R., Plastini, M.J., and Moraes, C.T. (2018). The mitochondrial DNA polymerase gamma degrades linear DNA fragments precluding the formation of deletions. Nat Commun 9, 2491. 10.1038/s41467-018-04895-1.

26. Fu, Y., Sacco, O., DeBitetto, E., Kanshin, E., Ueberheide, B., and Sfeir, A. (2023). Mitochondrial DNA breaks activate an integrated stress response to reestablish homeostasis. Molecular cell 83, 3740–3753 e3749. 10.1016/j.molcel.2023.09.026.

27. Peeva, V., Blei, D., Trombly, G., Corsi, S., Szukszto, M.J., Rebelo-Guiomar, P., Gammage, P.A., Kudin, A.P., Becker, C., Altmuller, J., et al. (2018). Linear mitochondrial DNA is rapidly degraded by components of the replication machinery. Nat Commun 9, 1727. 10.1038/s41467-018-04131-w.

28. Mimitou, E.P., Lareau, C.A., Chen, K.Y., Zorzetto-Fernandes, A.L., Hao, Y., Takeshima, Y., Luo, W., Huang, T.S., Yeung, B.Z., Papalexi, E., et al. (2021). Scalable, multimodal profiling of chromatin accessibility, gene expression and protein levels in single cells. Nat Biotechnol 39, 1246–1258. 10.1038/s41587-021-00927-2.

29. Malyarchuk, S., Wright, D., Castore, R., Klepper, E., Weiss, B., Doherty, A.J., and Harrison, L. (2007). Expression of Mycobacterium tuberculosis Ku and Ligase D in Escherichia coli results in RecA and RecB-independent DNA end-joining at regions of microhomology. DNA Repair (Amst) 6, 1413–1424. 10.1016/j.dnarep.2007.04.004.

30. Mullin, N.K., Voigt, A.P., Flamme-Wiese, M.J., Liu, X., Riker, M.J., Varzavand, K., Stone, E.M., Tucker, B.A., and Mullins, R.F. (2023). Multimodal single-cell analysis of nonrandom heteroplasmy distribution in human retinal mitochondrial disease. JCI Insight 8. 10.1172/jci.insight.165937.

31. Bayona-Bafaluy, M.P., Blits, B., Battersby, B.J., Shoubridge, E.A., and Moraes, C.T. (2005). Rapid directional shift of mitochondrial DNA heteroplasmy in animal tissues by a mitochondrially targeted restriction endonuclease. Proceedings of the National Academy of Sciences of the United States of America 102, 14392–14397. 10.1073/pnas.0502896102.

32. Bacman, S.R., Williams, S.L., and Moraes, C.T. (2009). Intra- and inter-molecular recombination of mitochondrial DNA after in vivo induction of multiple double-strand breaks. Nucleic Acids Res 37, 4218–4226. 10.1093/nar/gkp348.

33. Samuels, D.C., Schon, E.A., and Chinnery, P.F. (2004). Two direct repeats cause most human mtDNA deletions. Trends Genet 20, 393–398. 10.1016/j.tig.2004.07.003.

34. Basu, S., Xie, X., Uhler, J.P., Hedberg-Oldfors, C., Milenkovic, D., Baris, O.R., Kimoloi, S., Matic, S., Stewart, J.B., Larsson, N.G., et al. (2020). Accurate mapping of mitochondrial DNA deletions and duplications using deep sequencing. PLoS genetics 16, e1009242. 10.1371/journal.pgen.1009242.

35. Gonzalez, C.D., Nissanka, N., Van Booven, D., Griswold, A.J., and Moraes, C.T. (2024). Absence of both MGME1 and POLG EXO abolishes mtDNA whereas absence of either creates unique mtDNA duplications. J Biol Chem 300, 107128. 10.1016/j.jbc.2024.107128.

36. Fontana, G.A., and Gahlon, H.L. (2020). Mechanisms of replication and repair in mitochondrial DNA deletion formation. Nucleic Acids Res 48, 11244–11258. 10.1093/nar/gkaa804.

37. Hurley, J.B. (2021). Retina Metabolism and Metabolism in the Pigmented Epithelium: A Busy Intersection. Annu Rev Vis Sci 7, 665–692. 10.1146/annurev-vision-100419-115156.

38. King, M.P., and Attardi, G. (1989). Human cells lacking mtDNA: repopulation with exogenous mitochondria by complementation. Science 246, 500–503. 10.1126/science.2814477.

39. Birsoy, K., Wang, T., Chen, W.W., Freinkman, E., Abu-Remaileh, M., and Sabatini, D.M. (2015). An Essential Role of the Mitochondrial Electron Transport Chain in Cell Proliferation Is to Enable Aspartate Synthesis. Cell 162, 540–551. 10.1016/j.cell.2015.07.016.

40. Sullivan, L.B., Gui, D.Y., Hosios, A.M., Bush, L.N., Freinkman, E., and Vander Heiden, M.G. (2015). Supporting Aspartate Biosynthesis Is an Essential Function of Respiration in Proliferating Cells. Cell 162, 552–563. 10.1016/j.cell.2015.07.017.

41. Lareau, C.A., Ludwig, L.S., Muus, C., Gohil, S.H., Zhao, T., Chiang, Z., Pelka, K., Verboon, J.M., Luo, W., Christian, E., et al. (2021). Massively parallel single-cell mitochondrial DNA genotyping and chromatin profiling. Nat Biotechnol 39, 451–461. 10.1038/s41587-020-0645-6.

42. Virbasius, C.A., Virbasius, J.V., and Scarpulla, R.C. (1993). NRF-1, an activator involved in nuclear-mitochondrial interactions, utilizes a new DNA-binding domain conserved in a family of developmental regulators. Genes & development 7, 2431–2445. 10.1101/gad.7.12a.2431.

43. Wu, Z., Puigserver, P., Andersson, U., Zhang, C., Adelmant, G., Mootha, V., Troy, A., Cinti, S., Lowell, B., Scarpulla, R.C., and Spiegelman, B.M. (1999). Mechanisms controlling mitochondrial biogenesis and respiration through the thermogenic coactivator PGC-1. Cell 98, 115–124. 10.1016/S0092-8674(00)80611-X.

44. Corral-Debrinski, M., Horton, T., Lott, M.T., Shoffner, J.M., Beal, M.F., and Wallace, D.C. (1992). Mitochondrial DNA deletions in human brain: regional variability and increase with advanced age. Nat Genet 2, 324–329. 10.1038/ng1292-324.

45. Taylor, R.W., and Turnbull, D.M. (2005). Mitochondrial DNA mutations in human disease. Nat Rev Genet 6, 389–402. 10.1038/nrg1606.

46. Rossignol, R., Faustin, B., Rocher, C., Malgat, M., Mazat, J.P., and Letellier, T. (2003). Mitochondrial threshold effects. Biochem J 370, 751–762. 10.1042/BJ20021594.

47. Grady, J.P., Pickett, S.J., Ng, Y.S., Alston, C.L., Blakely, E.L., Hardy, S.A., Feeney, C.L., Bright, A.A., Schaefer, A.M., Gorman, G.S., et al. (2018). mtDNA heteroplasmy level and copy number indicate disease burden in m.3243A>G mitochondrial disease. EMBO Mol Med 10. 10.15252/emmm.201708262.

48. Hammans, S.R., Sweeney, M.G., Brockington, M., Lennox, G.G., Lawton, N.F., Kennedy, C.R., Morgan-Hughes, J.A., and Harding, A.E. (1993). The mitochondrial DNA transfer RNA(Lys)A-->G(8344) mutation and the syndrome of myoclonic epilepsy with ragged red fibres (MERRF). Relationship of clinical phenotype to proportion of mutant mitochondrial DNA. Brain 116 ( *Pt 3*), 617–632. 10.1093/brain/116.3.617.

49. Filograna, R., Mennuni, M., Alsina, D., and Larsson, N.G. (2021). Mitochondrial DNA copy number in human disease: the more the better? FEBS Lett 595, 976–1002. 10.1002/1873-3468.14021.

50. Lax, N.Z., Gorman, G.S., and Turnbull, D.M. (2017). Review: Central nervous system involvement in mitochondrial disease. Neuropathol Appl Neurobiol 43, 102–118. 10.1111/nan.12333.

51. Soto, I., Couvillion, M., Hansen, K.G., McShane, E., Moran, J.C., Barrientos, A., and Churchman, L.S. (2022). Balanced mitochondrial and cytosolic translatomes underlie the biogenesis of human respiratory complexes. Genome Biol 23, 170. 10.1186/s13059-022-02732-9.

52. Picard, M., Zhang, J., Hancock, S., Derbeneva, O., Golhar, R., Golik, P., O’Hearn, S., Levy, S., Potluri, P., Lvova, M., et al. (2014). Progressive increase in mtDNA 3243A>G heteroplasmy causes abrupt transcriptional reprogramming. Proceedings of the National Academy of Sciences of the United States of America 111, E4033–4042. 10.1073/pnas.1414028111.

53. McShane, E., Couvillion, M., Ietswaart, R., Prakash, G., Smalec, B.M., Soto, I., Baxter-Koenigs, A.R., Choquet, K., and Churchman, L.S. (2024). A kinetic dichotomy between mitochondrial and nuclear gene expression processes. Molecular cell 84, 1541–1555 e1511. 10.1016/j.molcel.2024.02.028.

54. Moran, J.C., Brivanlou, A., Brischigliaro, M., Fontanesi, F., Rouskin, S., and Barrientos, A. (2024). The human mitochondrial mRNA structurome reveals mechanisms of gene expression. Science 385, eadm9238. 10.1126/science.adm9238.

55. Debray, F.G., Morin, C., Janvier, A., Villeneuve, J., Maranda, B., Laframboise, R., Lacroix, J., Decarie, J.C., Robitaille, Y., Lambert, M., et al. (2011). LRPPRC mutations cause a phenotypically distinct form of Leigh syndrome with cytochrome c oxidase deficiency. J Med Genet 48, 183–189. 10.1136/jmg.2010.081976.

56. Kotrys, A.V., Durham, T.J., Guo, X.A., Vantaku, V.R., Parangi, S., and Mootha, V.K. (2024). Single-cell analysis reveals context-dependent, cell-level selection of mtDNA. Nature 629, 458–466. 10.1038/s41586-024-07332-0.

57. Forsstrom, S., Jackson, C.B., Carroll, C.J., Kuronen, M., Pirinen, E., Pradhan, S., Marmyleva, A., Auranen, M., Kleine, I.M., Khan, N.A., et al. (2019). Fibroblast Growth Factor 21 Drives Dynamics of Local and Systemic Stress Responses in Mitochondrial Myopathy with mtDNA Deletions. Cell Metab 30, 1040–1054 e1047. 10.1016/j.cmet.2019.08.019.

58. Khan, N.A., Nikkanen, J., Yatsuga, S., Jackson, C., Wang, L., Pradhan, S., Kivela, R., Pessia, A., Velagapudi, V., and Suomalainen, A. (2017). mTORC1 Regulates Mitochondrial Integrated Stress Response and Mitochondrial Myopathy Progression. Cell Metab 26, 419–428 e415. 10.1016/j.cmet.2017.07.007.

59. Komuro, A., Masuda, Y., Kobayashi, K., Babbitt, R., Gunel, M., Flavell, R.A., and Marchesi, V.T. (2004). The AHNAKs are a class of giant propeller-like proteins that associate with calcium channel proteins of cardiomyocytes and other cells. Proceedings of the National Academy of Sciences of the United States of America 101, 4053–4058. 10.1073/pnas.0308619101.

60. Mbaya, E., Oules, B., Caspersen, C., Tacine, R., Massinet, H., Pennuto, M., Chretien, D., Munnich, A., Rotig, A., Rizzuto, R., et al. (2010). Calcium signalling-dependent mitochondrial dysfunction and bioenergetics regulation in respiratory chain Complex II deficiency. Cell Death Differ 17, 1855–1866. 10.1038/cdd.2010.51.

61. Moudy, A.M., Handran, S.D., Goldberg, M.P., Ruffin, N., Karl, I., Kranz-Eble, P., DeVivo, D.C., and Rothman, S.M. (1995). Abnormal calcium homeostasis and mitochondrial polarization in a human encephalomyopathy. Proceedings of the National Academy of Sciences of the United States of America 92, 729–733. 10.1073/pnas.92.3.729.

62. Brini, M., Pinton, P., King, M.P., Davidson, M., Schon, E.A., and Rizzuto, R. (1999). A calcium signaling defect in the pathogenesis of a mitochondrial DNA inherited oxidative phosphorylation deficiency. Nat Med 5, 951–954. 10.1038/11396.

63. Li, M., Zhong, A., Wu, Y., Sidharta, M., Beaury, M., Zhao, X., Studer, L., and Zhou, T. (2022). Transient inhibition of p53 enhances prime editing and cytosine base-editing efficiencies in human pluripotent stem cells. Nat Commun 13, 6354. 10.1038/s41467-022-34045-7.

64. Ciceri, G., Baggiolini, A., Cho, H.S., Kshirsagar, M., Benito-Kwiecinski, S., Walsh, R.M., Aromolaran, K.A., Gonzalez-Hernandez, A.J., Munguba, H., Koo, S.Y., et al. (2024). An epigenetic barrier sets the timing of human neuronal maturation. Nature 626, 881–890. 10.1038/s41586-023-06984-8.

65. Zhou, T., Benda, C., Dunzinger, S., Huang, Y., Ho, J.C., Yang, J., Wang, Y., Zhang, Y., Zhuang, Q., Li, Y., et al. (2012). Generation of human induced pluripotent stem cells from urine samples. Nat Protoc 7, 2080–2089. 10.1038/nprot.2012.115.

66. Moretton, A., Morel, F., Macao, B., Lachaume, P., Ishak, L., Lefebvre, M., Garreau-Balandier, I., Vernet, P., Falkenberg, M., and Farge, G. (2017). Selective mitochondrial DNA degradation following double-strand breaks. PloS one 12, e0176795. 10.1371/journal.pone.0176795.

67. Buenrostro, J.D., Giresi, P.G., Zaba, L.C., Chang, H.Y., and Greenleaf, W.J. (2013). Transposition of native chromatin for fast and sensitive epigenomic profiling of open chromatin, DNA-binding proteins and nucleosome position. Nature methods 10, 1213–1218. 10.1038/nmeth.2688.

68. Kim, M., Gorelick, A.N., Vazquez-Garcia, I., Williams, M.J., Salehi, S., Shi, H., Weiner, A.C., Ceglia, N., Funnell, T., Park, T., et al. (2024). Single-cell mtDNA dynamics in tumors is driven by coregulation of nuclear and mitochondrial genomes. Nat Genet 56, 889–899. 10.1038/s41588-024-01724-8.

69. Lieber, T., Jeedigunta, S.P., Palozzi, J.M., Lehmann, R., and Hurd, T.R. (2019). Mitochondrial fragmentation drives selective removal of deleterious mtDNA in the germline. Nature 570, 380–384. 10.1038/s41586-019-1213-4.

70. Stoeckius, M., Zheng, S., Houck-Loomis, B., Hao, S., Yeung, B.Z., Mauck, W.M., 3rd, Smibert, P., and Satija, R. (2018). Cell Hashing with barcoded antibodies enables multiplexing and doublet detection for single cell genomics. Genome Biol 19, 224. 10.1186/s13059-018-1603-1.

71. Lareau, C.A., Liu, V., Muus, C., Praktiknjo, S.D., Nitsch, L., Kautz, P., Sandor, K., Yin, Y., Gutierrez, J.C., Pelka, K., et al. (2023). Mitochondrial single-cell ATAC-seq for high-throughput multi-omic detection of mitochondrial genotypes and chromatin accessibility. Nature protocols 18, 1416–1440. 10.1038/s41596-022-00795-3.

72. Persad, S., Choo, Z.N., Dien, C., Sohail, N., Masilionis, I., Chaligne, R., Nawy, T., Brown, C.C., Sharma, R., Pe’er, I., et al. (2023). SEACells infers transcriptional and epigenomic cellular states from single-cell genomics data. Nat Biotechnol 41, 1746–1757. 10.1038/s41587-023-01716-9.

73. van Dijk, D., Sharma, R., Nainys, J., Yim, K., Kathail, P., Carr, A.J., Burdziak, C., Moon, K.R., Chaffer, C.L., Pattabiraman, D., et al. (2018). Recovering Gene Interactions from Single-Cell Data Using Data Diffusion. Cell 174, 716–729 e727. 10.1016/j.cell.2018.05.061.

74. Chen, J., Lopez-Moyado, I.F., Seo, H., Lio, C.J., Hempleman, L.J., Sekiya, T., Yoshimura, A., Scott-Browne, J.P., and Rao, A. (2019). NR4A transcription factors limit CAR T cell function in solid tumours. Nature 567, 530–534. 10.1038/s41586-019-0985-x.

75. Heumos, L., Schaar, A.C., Lance, C., Litinetskaya, A., Drost, F., Zappia, L., Lucken, M.D., Strobl, D.C., Henao, J., Curion, F., et al. (2023). Best practices for single-cell analysis across modalities. Nat Rev Genet 24, 550–572. 10.1038/s41576-023-00586-w.

76. Morris, J.A., Caragine, C., Daniloski, Z., Domingo, J., Barry, T., Lu, L., Davis, K., Ziosi, M., Glinos, D.A., Hao, S., et al. (2023). Discovery of target genes and pathways at GWAS loci by pooled single-cell CRISPR screens. Science 380, eadh7699. 10.1126/science.adh7699.

77. Ganel, L., Chen, L., Christ, R., Vangipurapu, J., Young, E., Das, I., Kanchi, K., Larson, D., Regier, A., Abel, H., et al. (2021). Mitochondrial genome copy number measured by DNA sequencing in human blood is strongly associated with metabolic traits via cell-type composition differences. Hum Genomics 15, 34. 10.1186/s40246-021-00335-2.

78. Zhang, P., Lehmann, B.D., Samuels, D.C., Zhao, S., Zhao, Y.Y., Shyr, Y., and Guo, Y. (2017). Estimating relative mitochondrial DNA copy number using high throughput sequencing data. Genomics 109, 457–462. 10.1016/j.ygeno.2017.07.002.

